# Polymer-like model to study the dynamics of dynamin filaments on deformable membrane tubes

**DOI:** 10.1101/686873

**Authors:** Jeffrey K. Noel, Frank Noé, Oliver Daumke, Alexander S. Mikhailov

## Abstract

Peripheral membrane proteins with intrinsic curvature can act both as sensors of membrane curvature and shape modulators of the underlying membranes. A well-studied example of such proteins is the mechano-chemical GTPase dynamin that assembles into helical filaments around membrane tubes and catalyzes their scission in a GTPase-dependent manner. It is known that the dynamin coat alone, without GTP, can constrict membrane tubes to radii of about 10 nanometers, indicating that the intrinsic shape and elasticity of dynamin filaments should play an important role in membrane remodeling. However, molecular and dynamic understanding of the process is lacking. Here, we develop a dynamical polymer-chain model for a helical elastic filament bound on a deformable membrane tube of conserved mass, accounting for thermal fluctuations in the filament and lipid flows in the membrane. The model is based on a locally-cylindrical helix approximation for dynamin. We obtain the elastic parameters of the dynamin filament by molecular dynamics simulations of its tetrameric building block and also from coarse-grained structure-based simulations of a 17-dimer filament. The results show that the stiffness of dynamin is comparable to that of the membrane. We determine equilibrium shapes of the filament and the membrane, and find that mostly the pitch of the filament, not its radius, is sensitive to variations in membrane tension and stiffness. The close correspondence between experimental estimates of the inner tube radius and those predicted by the model suggests that dynamin’s “stalk” region is responsible for its GTP-independent membrane-shaping ability. The model paves the way for future mesoscopic modeling of dynamin with explicit motor function.

## 1 Introduction

The shape of biological membranes is controlled by proteins that stabilize and/or generate membrane curvature [1]. During the well-studied process of clathrin-mediated endocytosis, protein scaffolds with a curved membrane binding surface are recruited to clathrin-coated pits and superimpose their curvature to the underlying membrane [2]. As a paradigm for this mechanism, Bin/amphiphysin/Rvs(BAR)-domain containing proteins form dimeric modules of various curvatures that drive different stages of endocytosis. Such membrane reshaping processes are accomplished passively, i.e. without the input of energy. Conversely, the final stage of endocytosis is catalyzed by the mechano-chemical enzyme dynamin that assembles at the endocytic neck and uses the energy of GTP hydrolysis for scission. Dynamin can be considered as a chimeric protein. While it acts as a polymeric structural scaffold that coils around the membrane tube and stabilizes membrane curvature, it also works as a motor protein that generates the active forces required to perform membrane scission. Understanding the mechanical properties of the dynamin filament is crucial to uncover how it transmits motor force toward the constriction and eventual scission of the underlying membrane template.

Dynamin’s structure and function in membrane scission have been extensively experimentally studied, as summarized in the detailed review [3]. Although endocytosis involves a complex orchestration of dozens of different proteins [4], it is remarkable that, in the presence of GTP, dynamin alone is sufficient to constrict the membrane, create torque, and perform scission *in vitro* [5, 6]. Through a combination of cryo-electron microscopy [7, 8] and X-ray crystallography [9, 10, 11] the structure of the dynamin oligomer on a membrane template has been elucidated (Fig. 1). Two-fold symmetric dimers of dynamin are formed by a central interface in the stalk. These dimers further assemble into tetramers and polymers through two additional stalk interfaces (the tetramer interface [11]). The resulting stalk filament has an intrinsic helical shape and acts as a scaffold that stabilizes membrane curvature. The pleckstrin homology (PH) domains protrude from the filament toward the membrane and mediate binding to the lipid surface [12, 13, 14]. Opposite the PH domains are the motor domains which are composed of a GTPase domain (G-domain) and a bundle signaling element (BSE). GTP-induced dimerization between G-domains of neighboring turns triggers GTP hydrolysis and results in a conformational change of the BSE relative to the G-domain [6, 15]. The resulting powerstroke is thought to pull adjacent turns of the filament along each other, leading to the further constriction of the filament and of the underlying membrane. Progress in high-speed atomic force microscopy (HS-AFM) recently allowed direct observation of membrane constriction and fission by dynamin at sub-10 nanometer resolution in real time [16, 17].

**Figure 1:**
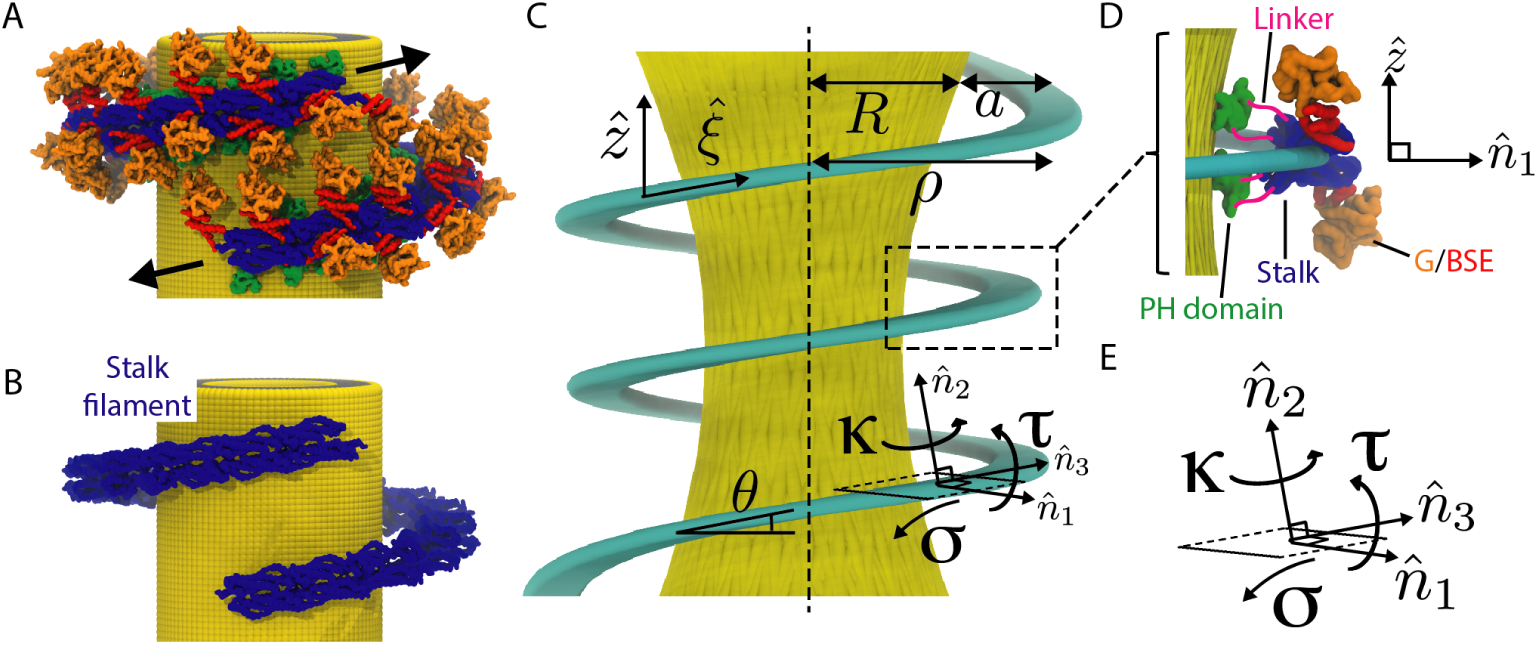
Geometry of the dynamin filament on a membrane tube. A) Cartoon of a functional dynamin filament consistent with X-ray crystallography and cryo-EM. In presence of GTP, motor domains (orange/red) mediate interactions between turns and produce torque (arrows). B) The stalk filament corresponds to the modeled elastic ribbon. C) The coordinates and variables employed in the continuous elastic description. Membrane tube in yellow and elastic ribbon in cyan. D) Apolar symmetry of the dynamin dimer. The PH domains (green) have affinity for the membrane and are located next to the membrane tube; they are connected by flexible linkers (pink) to the stalk domains. The motor domains point away from the membrane. The filament is formed by connections between the stalk domains (blue). E) The local material frame for the filament. The normal curvature *κ*, twist *τ*, and geodesic curvature *σ* correspond to rotations about the axes 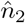, 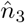, and 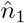, respectively.

Although dynamin’s helical geometry and membrane template hinder direct measurement of the elastic parameters, biophysical measurements have shed some light on these factors. Early experiments provided evidence that the stalk filament’s intrinsic curvature confers dynamin’s ability to tubulate flat membranes even in the absence of motor activity [5]. More recently [18], the analysis of an optical tweezer setup using the elastic theory of membrane tubes [19, 20] suggested that the spontaneous curvature was similar to the curvature observed in crystal/cryo-EM structures. Studies have also highlighted the role of membrane tension in accelerating membrane scission [21]. An important aspect of the conjectured constriction mechanism of scission is the flexibility of the dynamin scaffold. As a constricting scaffold (Fig. 1A), the dynamin filament must not be so prohibitively stiff that it prevents its own tightening. On the other hand, dynamin must be sufficiently stiff so that it can spontaneously (i.e. in the absence of GTP) remodel membranes into tight tubes of 10-20 nanometers in diameter [5, 18].

Dynamin has been theoretically investigated at different levels, ranging from all-atom molecular dynamics (MD) simulations [9, 11] to coarse-grained modeling [22, 23], to mesoscopic continuum descriptions for the filament and the membrane. Our focus is on the latter kind of models.

The present paper is the first step toward our aim to develop a platform for systematic stochastic dynamical simulations of dynamin filaments on deformable membrane tubes. While eventually the GTP-hydrolysis effects of motor activity and dimerization of G-domains can be incorporated into the developed model, the present publication shall only deal with the situation in absence of GTP. At this stage, processes of transition to an equilibrium state of the membrane and the filament, and thermal fluctuations around this state will be considered and analyzed.

As originally proposed by M. Kozlov [24], the stalk filament of dynamin can be modeled as a thin and narrow elastic ribbon with spontaneous normal curvature and twist. The important elastic parameters of the stalk filament are its stiffness moduli with respect to variations in the normal and geodesic curvature and the twist; they describe how rigid the filament is. The relationship between the elastic coefficients of the filament and those of the membrane determines whether the filament would be more stiff or more soft than the membrane tube. While elastic parameters of lipid bilayers are experimentally well-known, there are no measurements of such parameters for dynamin. In a previous work by some of the authors [9], the free energy surface of a modeled tetramer was computed, and it predicted that the stalk filament should have significant intrinsic curvature and twist. However, the filament’s elastic parameters were not yet quantified. In absence of this knowledge, different modeling limits have been phenomenologically considered, assuming that either the filament [24, 25, 18, 26, 27] or the membrane tube [28] were predominantly stiff.

In our study, the elastic parameters of the stalk filament are determined from specially designed microscopic simulations. By performing all-atom MD simulations for a tetramer and analyzing their results, we obtain the spontaneous curvature and twist, together with the stiffness constants for variations of the normal curvature and the twist, and the stretch modulus for dynamin. Additional coarse-grained simulations of a 17-dimer oligomer were performed to determine the stiffness constant with respect to variations of the geodesic curvature. We find that the elasticity of dynamin is well tuned to that of the membrane, so that both the filament can deform the membrane and the membrane can significantly act back on it. This property, that has probably emerged in the evolution of dynamin, is related to the presence of a special normal mode in the protein. This soft mode reflects flexibility in the tetramer interface that explains how the stalk filament can adapt to varying curvature of the membrane tube.

At the next stage, interactions between the stalk filament and the membrane are introduced. Microscopically, such interactions arise because dimers forming the stalk are connected by flexible linkers to PH domains that attach upon the membrane. In our model, coupling of the stalk filament to the membrane through flexible linkers is effectively taken into account by a strong interaction potential ensuring that the local radius of the filament coincides with the local outer radius of the membrane tube.

Several modeling assumptions are made by us. Only axially-symmetric membrane tubes with the radius slowly varying along the tube will be considered, so that, locally, the membrane has a cylindrical shape. Similarly, we assume that the filament locally represents a cylindrical helix, with a radius and a pitch that slowly vary along it. Moreover, in the present study, the length of the filament is not allowed to change, thus omitting polymerization effects. Formally, this limits our analysis to the situations near to the dynamin nucleation point.

Using continuum theory, we determine approximate equilibrium shapes of the filament on a membrane tube. We find that the lumen radius of a dynamin-covered membrane tube varies only a little, 10-12 nanometers, over a range of physiologically-relevant stiffness and tension values of the membrane. On the other hand, it is found that the membrane can strongly compress the filament in the axial direction. This causes the pitch of the filament to be sensitive to the membrane parameters, undergoing a transition from a compressed phase to an expanded phase as the membrane tension is increased.

Despite these new results, the continuum model essentially represents an intermediate stage in the construction of our stochastic polymer-like model for dynamin filaments on deformable membrane tubes. In this model, evolution equations are formulated in terms of a polymer-like description where a filament is viewed as a chain of interacting beads. Each bead corresponds to one dynamin dimer. This stretchable polymer has spontaneous curvature and twist, tending to form a helix. Overdamped stochastic dynamics of beads, governed by Langevin equations with thermal noise, is assumed.

For the membrane, a continuous evolution equation is first derived by using the conservation law for its mass. This equation explicitly takes into account hydrodynamical lipid flows in the membrane. Interactions between the membrane and the beads are then introduced: the filament radially compresses the membrane and can also drag it by viscous friction in the axial direction. After that, discretization is performed, i.e. the tube is divided into a set of thin discs.

Using the polymer model, numerical simulations have been undertaken. Thus, we could determine equilibrium shapes for filaments of various sizes, assuming different tension and bending rigidity of the membrane. In such simulations, the effects of axial filament compression by the membrane could be directly seen and the results of the approximate analytical theory could be verified. In the simulations, the helical filament is characterized by axially-varying curvature and twist, with the gradients getting stronger near the free ends of it. Therefore, we could also check then the validity of our model assumption that the radius of the filament (and the membrane tube) and the pitch of the filament vary only slowly along it. Because the model includes thermal noise for the beads, thermal fluctuations in the local shapes could also be explored.

For clarity reasons, we have not followed the actual presentation structure in our brief summary of the contents. Below, the complete model construction is first undertaken and then the results of analytical and numerical investigations are described.

## 2 Model Development

### 2.1 The continuum elastic description

Our primary goal is to develop a polymer chain model of the dynamin filament that is amenable to Brownian dynamics simulations. We begin by formulating a continuum elastic description that will serve as the theoretical ground for a polymer chain model. Additionally, in the Results, the continuum model will be used to provide an analytical explanation for the filament shape changes seen in numerical simulations.

The geometry of the dynamin filament on a straight and axially-symmetric membrane surface (henceforth referred to as the membrane tube) is illustrated in Fig. 1. The structural part of the dynamin polymer is referred to as the stalk filament (Fig. 1B). The filament is attached to the membrane tube by flexible linkers that connect stalk domains to PH domains that attach upon the membrane. First, we consider the deformation energy of the filament apart from the membrane. After that, the elastic energy of the membrane is considered and interactions between the membrane and the filament are introduced.

#### 2.1.1 Elastic energy of the filament

Dynamin oligomerization creates a filament that naturally takes on a helical shape, which, under special conditions, can even be observed in the absence of the membrane [7]. As originally proposed by M. Kozlov [24], this filament can be modeled as a thin and narrow ribbon (Fig. 1C). Generally, such a ribbon is defined [29] by its center line and by local orientations of three mutually orthogonal unit vectors (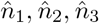) that form the material reference frame. The vectors 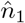 and 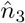 lie in the plane of the ribbon, while the vector 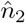 is orthogonal to it. The central line together with the set of material frames attached to it at every point represents a *Cosserat curve* [29]. Below, the width and the thickness of the ribbon do not enter into the description and, essentially, the stalk filament is modeled as such an oriented curve. The normal curvature *κ*, twist *τ*, and geodesic curvature *σ* correspond to rotations about the axes 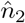, 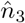, and 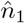, respectively, when moving along the central line. If *ξ* is the arc length along this line, one has [29]

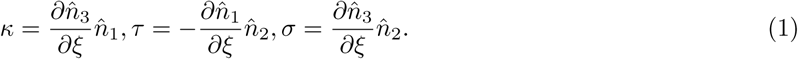

Different choices for the local orientations of vectors (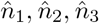) with respect to the membrane surface can be made. It is often assumed that the ribbon stays orthogonal to the membrane, so that the vector 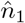 coincides with the local normal vector of the membrane. In the recent publication [30], it was however pointed out that the ribbon can be also tilted with respect to the membrane, so that the vector 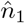 is no longer orthogonal to it. Additional tilt energy was included and its effects on equilibrium shapes of uniform filaments were discussed [30]. In our description, interactions between the filament and the membrane will be explicitly introduced below in Section 2.1.3 and then different assumptions about such interactions might be made.

It is known that the equilibrium shape of the dynamin filament is a uniform straight helix of some radius *ρ*_0_ and pitch *h*_0_. For the uniform helix, the vector 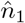 is orthogonal to the symmetry axis 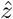. The geodesic curvature is vanishing for this helix, *σ* = 0. Its normal curvature and the twist are

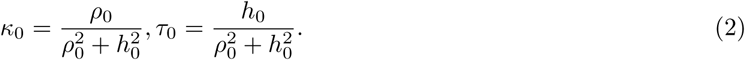

These intrinsic curvatures are related to the structure of the oligomerization interface and we refer to them as the *spontaneous* curvature and twist.

A deformation of the ribbon (as a Cosserat curve) will generally consist both of a deformation of its central line and a change in the orientations of the local material frames. Therefore, the ribbon will have normal and geodesic curvatures, and twist, that vary along its central line, *κ* = *κ*(*ξ*), *σ* = *σ*(*ξ*), *τ* = *τ*(*ξ*). These three properties characterize local deformations of the ribbon and the filament energy *E*_F_ should be expressed in terms of them. In the framework of the elasticity theory, the energy should be quadratic in deformations. Moreover, the minimum of this energy should correspond to the uniform helix *σ* = 0, *κ* = *κ*_0_, *τ* = *τ*_0_. Therefore, the deformation energy of the filament of length ℒ is [29]

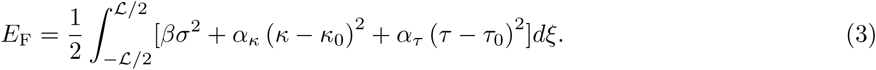

Here, *β, α*_*κ*_ and *α*_*τ*_ are the elastic moduli (i.e., the stiffness constants) with respect to the geodesic and normal curvatures, and the twist.

Deformed states of a ribbon generally correspond to quite complex shapes and even finding the equilibrium shapes of elastic curves constrained to various surfaces represents a difficult problem in differential geometry, see, e.g. [31]. Our aim is to develop a dynamic description for dynamin filaments interacting with deformable membranes. This description would have become too complicated and practically intractable if it were intended to correctly reproduce all possible evolving shapes. Therefore, some simplifications are to be made.

We shall consider only the filaments that locally have the cylindrical helix shape. Such helices will not be bent and will have a straight axial symmetry line 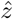. Their radius and pitch will be allowed to vary along the symmetry axis, so that *ρ* = *ρ*(*z*) and *h* = *h*(*z*). In our approximation it will be assumed that the radius is only slowly varied along the filament, so that it locally maintains a cylindrical shape. Namely, the condition (*dρ/dz*)^2^ *≪* 1 must be satisfied. Moreover, we shall assume that the orientations of the material frame vectors remain the same as for a uniform helix of the same radius and pitch. Particularly, the vector 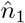 stays then orthogonal to the axis 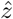.

The derivation of the locally-cylindrical approximation for the filament is provided in Supplementary Section 1. As we find, the normal curvature and the twist are approximately determined as

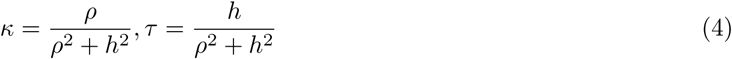

by the local radius, twist and pitch. The geodesic curvature depends on the gradients of these variables and, approximately, is

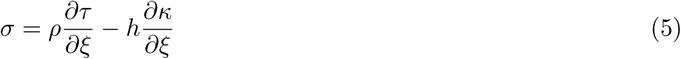

Using Eq. 5 and the inverse relationships

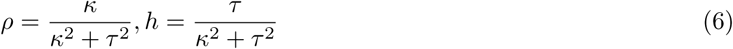

the elastic energy of the filament can be finally written as

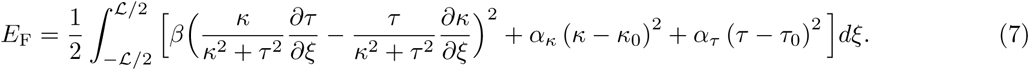

The first term, arising from the geodesic curvature, leads to coupling of the normal curvature and the twist between the elements of the filament. Numerical values of spontaneous curvature and twist, as well as the three elastic moduli, will be estimated for dynamin in Section 4.1.

Instead of *h*, the structural pitch *p* = 2*πh* will often be used. Moreover, the local twist angle defined by

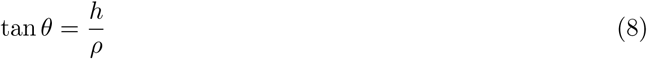

is introduced.

#### 2.1.2 Elastic energy of the membrane tube

Locally, the membrane surface is described by its two principal curvatures *c*_1_ and *c*_2_. According to Helfrich [32], the elastic energy of the membrane is given by

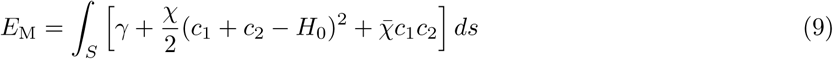

where *γ* is the surface tension coefficient, *χ* is the bending stiffness of the membrane, 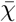 is the Gaussian curvature modulus, and *H*_0_ is the spontaneous curvature. The integration is performed along the entire surface *S* of the membrane. The spontaneous curvature of the free membrane will be taken to be zero. It can be noticed that terms of the form *c*_1_*c*_2_ do not generate any local forces for the considered axisymmetric geometry. Therefore, we will always drop such terms (see Supplementary Eq. 61 and also a general derivation in the review [33]).

Generally, the principal curvatures can depend both on the coordinate *z* along the tube and on the polar angle. To simplify the description, we shall however assume the axial symmetry of the tube. For such a membrane tube aligned along the *z*-axis and with slowly varying radius *R*(*z*), the first principal curvature can be chosen as the curvature in the horizontal cross-section (*x, y* plane) and the second curvature along the symmetry axis 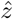. If the condition (*dR/dz*)^2^ *≪* 1 is satisfied, we approximately have

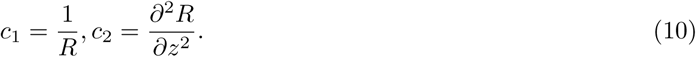

Moreover, we can then also approximately write 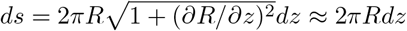.

Hence, in this locally-cylindrical approximation, the Helfrich elastic energy of a membrane tube of length *L* is

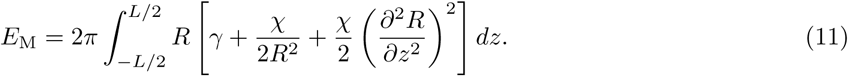

Physically, the membrane is a lipid bilayer and the radius *R* corresponds to the local distance from the tube axis to the middle surface of the bilayer.

Several methods exist for measuring the membrane bilayer elastic modulus *χ* (Eq. 9), these are reviewed in [34]. The parameter *χ* can vary depending on factors such as temperature, lipid composition and solvent. It has been demonstrated that the lipid composition of the clathrin-coated pit and neck is in flux during endocytosis [35], suggesting that there is likely a range of membrane rigidities encountered by dynamin during its function. A relevant in vitro study [18] of dynamin polymerization used membranes with a bending rigidity of *χ ≈* 16 k_B_T. In our study, the range of stiffnesses between this value and 24 k_B_T will be explored. Also important is the membrane tension *γ* because this sets the equilibrium radius of the membrane tube *R*_eq_ in the absence of protein [19]. This radius can be calculated by taking *∂E*_M_/*∂R* = 0 (Eq. 11):

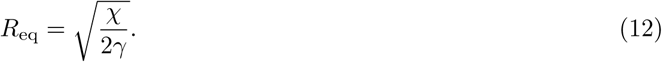

In this way, the tension *γ* can be used to tune the tube radius experimentally. During clathrin-mediated endocytosis, tension is present from the cell membrane and additional forces are created by actin polymerization [36, 4].

#### 2.1.3 Coupling between the filament and membrane

Finally, interactions between the stalk filament and the membrane are to be introduced. The stalk domains of dynamin are connected by linkers to the PH domains, which have affinity for phosphoinositide lipids in the membrane and, additionally, may insert a hydrophobic loop into the bilayer. The linkers (~ 20 amino acids in length at both the N- and C-terminus of the PH domain) are sufficiently flexible so that they cannot be resolved in X-ray crystal structures [9, 10, 11]. Additionally, the highest resolution cryo-EM map indicates random loop structures for them [8]. Further, the only structural evidence for a direct interaction interface between the PH domains and the stalks is an auto-inhibitory interface that is occluded in the stalk filament [11]. Therefore, in the context of the filament energy, the role of the PH domains seems to simply be to adhere through the linkers the filament to the membrane. According to this view, the flexible linkers cannot fix the orientation of the filament with respect to the PH domains that are free to independently adapt their own orientation to the local membrane shape (see also Fig. S1 in Supplementary Information).

Note that, in the recent publication [30], a different interaction mechanism with the membrane was chosen. It has been assumed that PH domains were rigidly attached to the stalk filament, therefore changing their orientations together with it with respect to the membrane. The basis for such assumption may have come from observations that PH domains are seen to rotate with respect to the membrane in a cryo-EM reconstruction of a super-constricted dynamin filament [37]. The developing tilt of PH domains led then, due to their interaction with the membrane, to an additional contribution in the energy. The involved effects on the shapes of the filaments were extensively discussed [30].

In contrast to this, in our model, coupling of the filament to the membrane via highly flexible linkers, as suggested by the above structural data, is chosen. Hence, the stalk filament will be allowed to have a local tilt with respect to the membrane, but there will be no additional energy associated with this tilt.

We do not explicitly define the linkers, but just assume that a harmonic interaction, keeping the center of the filament at a certain distance *a* from the middle of the lipid bilayer, is present. The value for *a* is taken from the cryo-EM structure of the “non-constricted state” [38] and *a* = 8.5 nanometers. Explicitly, the coupling interaction energy is

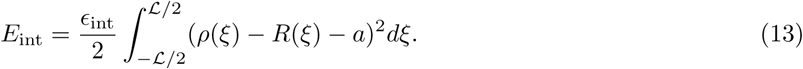

Here, *ρ*(*ξ*) and *R*(*ξ*) are both measured relative to the axis of the tube, and *R*(*ξ*) refers to the radius of the membrane tube at the position corresponding to the coordinate *ξ* along the filament. Formally, if the dependences of *κ* and *τ* on the coordinate *ξ* are known, the vertical position *z* of the membrane element with the coordinate *ξ* is

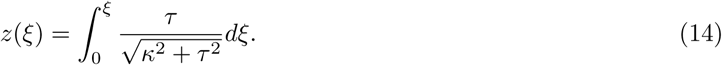

Therefore, *R*(*ξ*) = *R*(*z*(*ξ*)). In numerical simulations, the coefficient *ϵ*_int_ will be chosen large enough to ensure that significant deviations of *ρ* from *R* + *a* do not occur.

The total energy of the filament interacting with the deformable membrane tube in the continuum approximation is therefore

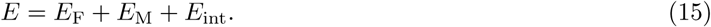

Note that a tilt energy could have been included into the model by introducing an additional energy term that depends on the scalar product of the vector that is normal to the membrane surface and the vector 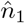 of the local material frame. Therefore, the orientational coupling mechanism of Ref.[30] may be incorporated as a model extension into our description too.

### 2.2 The discrete polymer model

The continuous elastic description for the filament/membrane system will be used in the Results to determine energies and uniform equilibrium states. While, in principle, it can be extended to also consider the dynamics of the filament and the membrane, this description gets then too complicated for practical use. Therefore, we further develop a discrete polymer-like model for the filament (and shall also introduce a discrete description for the membrane). This allows explicit mesoscale dynamical simulations for dynamin and the membrane where thermal fluctuations can be accounted too. Now, in the polymer model, stretching elasticity of the filament is additionally taken into account and repulsive excluded-volume interactions between the turns are introduced.

The dynamin filament is modeled by *N* dynamin beads, where each bead corresponds to a dynamin dimer (Fig. 2). The coordinates of bead *i* are 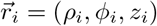 where *ρ*_*i*_ is the distance from the axis of the membrane, *φ*_*i*_ is the polar angle and *z*_*i*_ is the distance along the length of the tube. The membrane is modeled as a stack of *M* disks with the thickness of *d*_M_ = 4 nm, which is roughly the bilayer thickness and provides a natural minimum length for an elastic membrane description. Below, the energies of the polymer model are formulated by discretizing the continuum elastic energies. Then, the equations of motion for the polymer and the membrane are derived.

**Figure 2:**
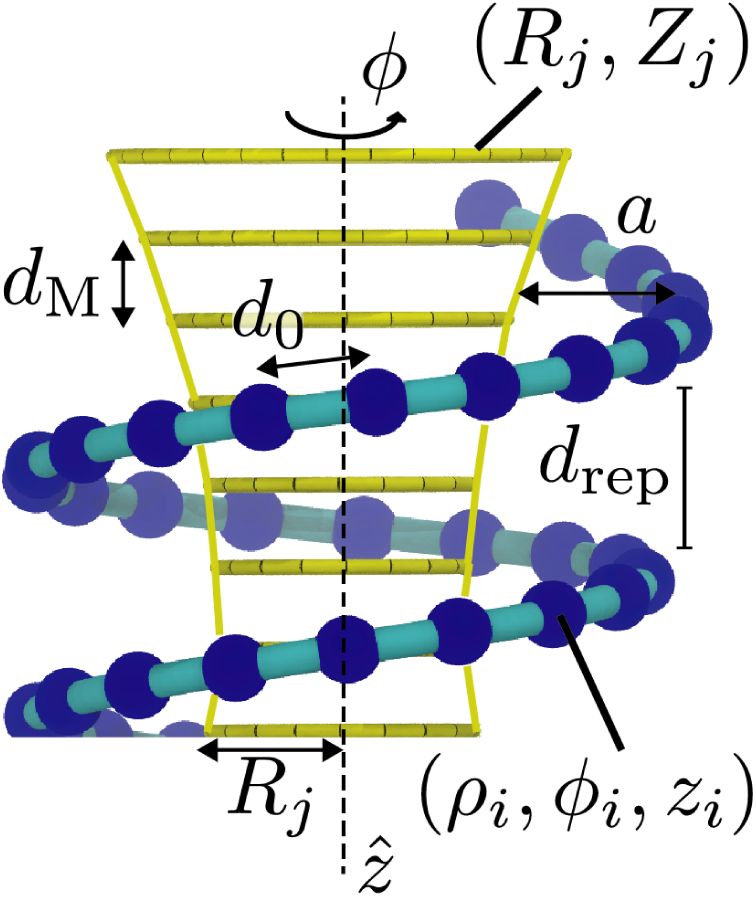
Definition of variables and parameters used in the polymer description.

#### 2.2.1 The discretization procedure

The energy of the polymeric filament is obtained by discretizing Eq. 7 into a sum over *N* polymer beads, with each term weighted by *d*_0_, the distance between successive dimers. Discretization of Eq. 7 involves determining the curvature *κ*_*i*_ and the twist *τ*_*i*_ at discrete positions 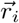. Using Eq. 4, we find

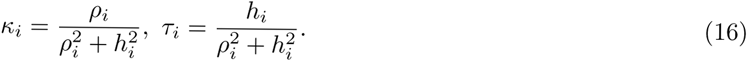

The coupling between the membrane and the filament (Eq. 13) enforces the filament to adopt the shape of a membrane tube with a slowly varying radius. Since the tube is aligned along the *z*-axis, the local pitch is given by *∂z/∂φ* and is expressed in terms of the filament coordinates as

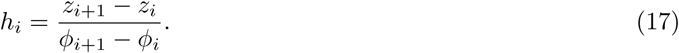

The geodesic curvature contains derivatives with respect to the filament coordinate *ξ*. These derivatives are discretized in the same manner as *h*_*i*_,

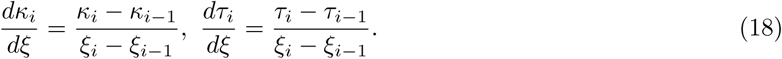

Note that the denominator is simply the distance between beads *ξ*_*i*_ − *ξ*_*i−*1_ ≡ *d*_0_.

While the filament is non-stretchable in the continuum approximation, the polymeric representation requires explicit interactions between the beads. In the polymer model, the connection is implemented by a harmonic pair interaction potential between the neighboring beads:

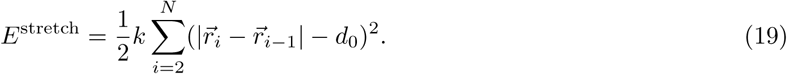

The filament stretching modulus *k* was obtained from the MD simulations using the distribution shown in Fig. 3D.

**Figure 3:**
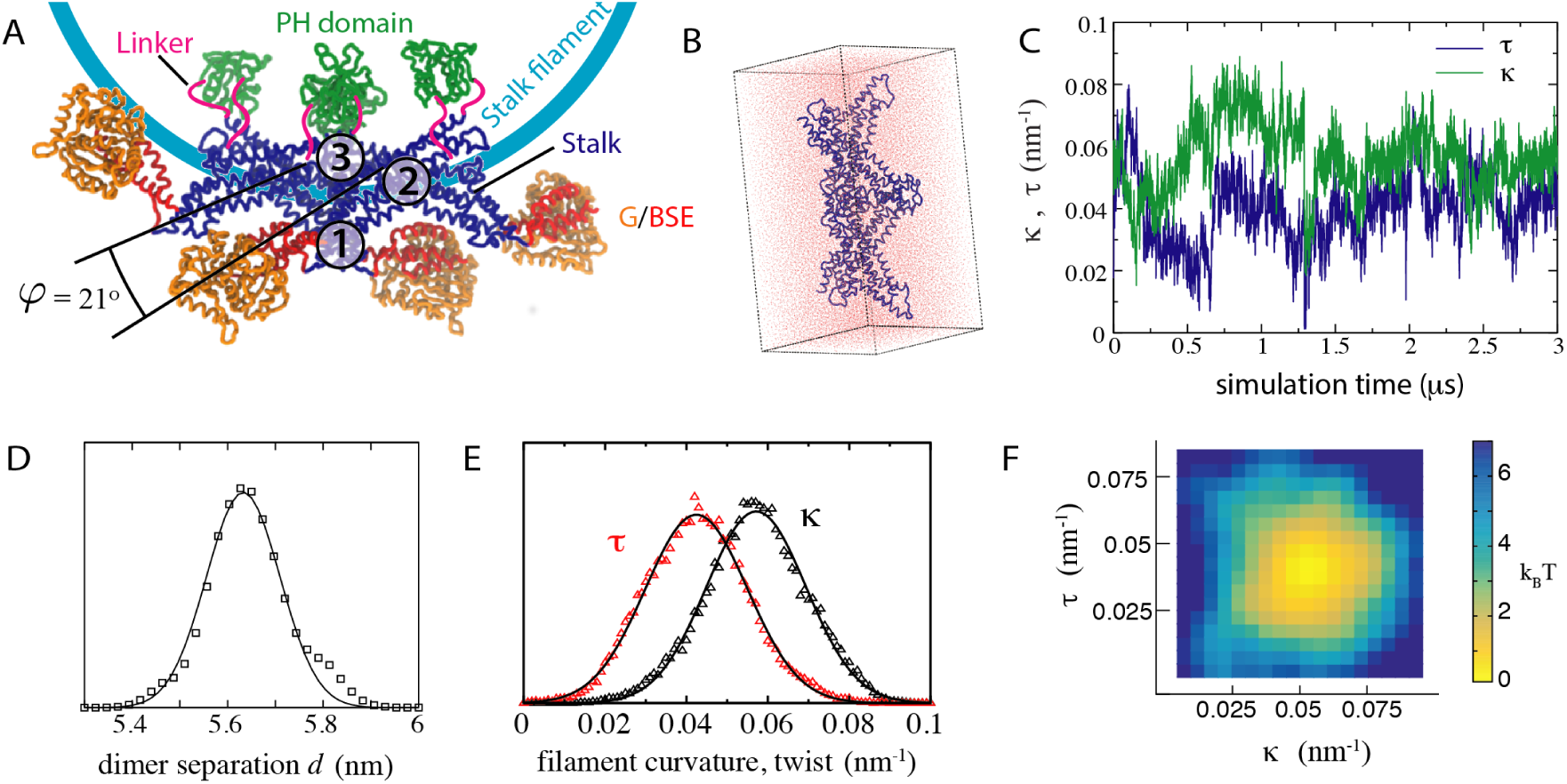
Estimating principal elastic parameters for the stalk tetramer from explicit-solvent molecular dynamics simulations. A) Architecture of the dynamin stalk tetramer. Interfaces 2 (dimer) and 1/3 (tetramer) are indicated. Here, the PH domain locations are not from X-ray crystallography, but are placed approximately as they appear in cryo-EM reconstructions. B) The stalk domains of the tetramer comprise the simulated system. C) Time dependence of curvature *κ* and twist *τ* over the explicit solvent trajectory. D) Distribution of distances *d* (squares) and its Gaussian fit (solid line); *d* is the distance between the centers of mass of the dimers. E) Distributions of *κ* (black triangles) and *τ* (red triangles), and their Gaussian fits (solid lines). F) Two-dimensional histogram of *κ* and *τ* represented as a potential of mean force.

Furthermore, excluded volume effects will be taken into account in the polymer model. They will determine how closely the neighboring turns can approach one another. To include such effects, we introduce a repulsive potential along the *z* direction:

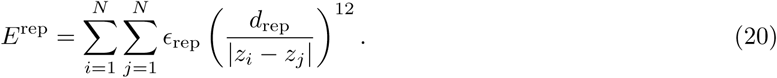

Here, *ϵ*_rep_ specifies the strength of repulsion and *d*_rep_ determines the repulsion length.

Taken all together, the energy of the dynamin polymer is given by

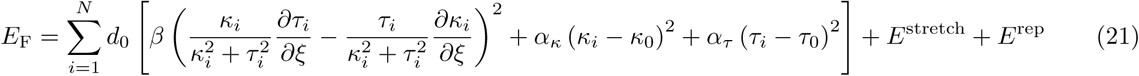

The discrete description for the membrane is derived from the axially-symmetric continuous Helfrich elastic membrane description, formulated above. After the discretization, the membrane tube consists of a stack of *M* disks of radii *R*_*i*_ centered along the *z*-axis at positions *Z*_*j*_ = −*L*/2 + *jd*_M_. The centers of the disks are fixed, but their radii can change. The dimensions of a bilayer provide a natural discretization for the membrane disks, *d*_M_ = 4 nm. The integral in Eq. 11 becomes a sum over the stack of disks

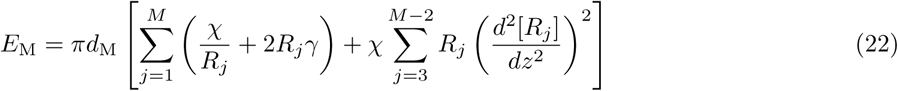

where a shorthand notation

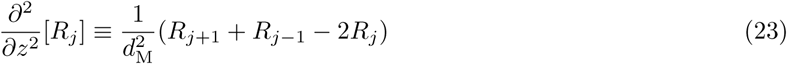

is used. We enforce that *∂*^2^/*∂z*^2^[*R*_*j*_] ≡ 0 at the membrane ends, *j* = 1, *M*.

After the discretization, the coupling energy between the membrane and the filament is

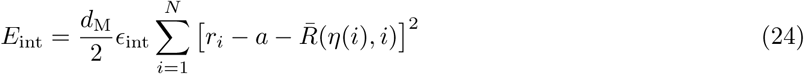

where the coefficient *ϵ*_int_ specifies the coupling strength. The role of this interaction is simply to keep filament beads at a constant distance *a* from the membrane. Here, we denote as *η*(*i*) a function that returns the index *j* of the membrane disk which includes the filament bead *i*, so that *Z*_*j*_ *< z*_*i*_ *< Z*_*j*+1_ where *z*_*i*_ is the coordinate of bead *i*. Furthermore, 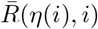 denotes a linear interpolation of the membrane radius between the nearest two membrane disks is used, to increase the approximation accuracy. Explicitly,

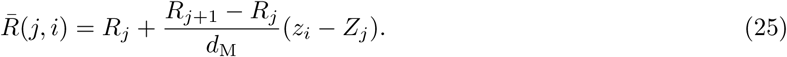

Adding all three components, the full potential energy *E* of the system is given by *E* = *E*_F_ + *E*_M_ + *E*_int_.

It should be noted that, while the energies have been obtained by discretization of the continuous elastic description, the discretization length is not chosen to provide an accurate quantitative approximation for the continuous case. Rather, the employed discretization length is chosen as the actual distance between dynamin dimers in the filament – allowing a direct interpretation of the beads as individual dimers. This representation is likely more appropriate than the continuum model since flexible connections between rigid dimers were suggested by the analysis in Section 4.1. In a fact, our polymer model constitutes an independent dynamical description, rather than a finite-difference scheme intended to accurately reproduce the continuous description of dynamin.

#### 2.2.2 Evolution equations for the filament

The motion of beads in the polymer is determined by forces applied to them. On the considered microscopic length scales, inertial effects are negligible and viscous friction effects prevail. There are two different sources of friction for dynamin. First, its dimers (corresponding to beads) move inside a viscous water solution. Second, the PH domains (see Fig. 1) are interacting with a lipid bilayer and move upon it. Because the viscosity of lipid bilayers is about 1000 times higher than that of water, we will assume that the dominant friction comes from interactions with the membrane.

Hence, the motion of bead *i* over the tube is described by two coupled evolution equations for its coordinates *φ*_*i*_ and *z*_*i*_:

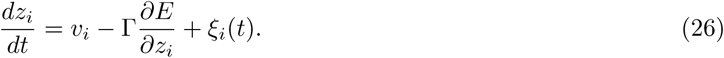

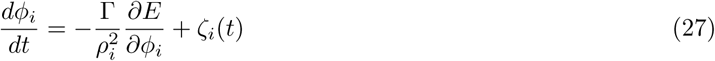

where Γ is taken to be the mobility of a single lipid within the membrane. The local lipid flow velocity *v*_*i*_ along the tube at the bead position *z* = *z*_*i*_ in the first equation accounts for the advection of beads by the membrane flow. Thermal fluctuations are included through random Gaussian thermal forces *ξ*_*i*_(*t*) and *ζ*_*i*_(*t*) with correlation functions

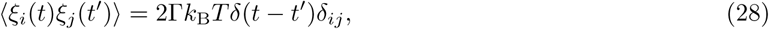

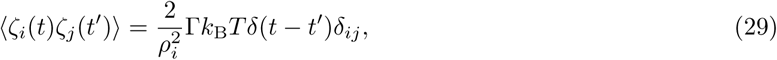

where *T* is the temperature and *k*_B_ is the Boltzmann constant.

Moreover, we need to formulate an evolution equation for the radius variable *r*_*i*_ of a bead. Because the filament is attached to the membrane, this variable should practically coincide with the local radius of the tube. The description is however simplified when it is allowed to slightly deviate from the tube radius, with a high energetic penalty described by the coupling energy *E*_int_ (Eq. 24). Thermal fluctuations can be neglected for this variable and the respective evolution equation will be

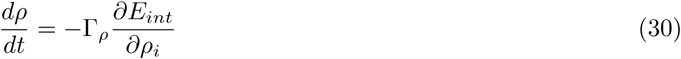

where Γ_*ρ*_ is a radial mobility. Because the coupling energy is proportional to the interaction strength parameter *ϵ*_int_ that should be large, but otherwise is arbitrary, we will make Γ_*ρ*_ = Γ by appropriate rescaling of the coefficient *ϵ*_int_.

#### 2.2.3 Evolution equation for the membrane

For the membrane tube, we first derive the evolution equation in the continuum description and then discretize it. Because lipid bilayers are practically incompressible, the total area of the membrane is approximately conserved. Hence, we can start with the continuity equation for the local area per unit length *s* of the membrane tube:

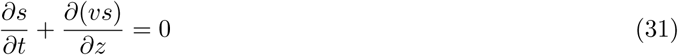

where *v* is the local lipid flow velocity. The tube area between *z* + ∆*z* and *z* is 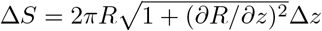 and therefore the area per unit length is 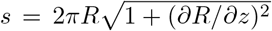. Assuming that the slope is sufficiently small, i.e. that (*dR/dz*)^2^ ≪ 1, we can approximately write *s* = 2*πR*. Hence, the continuity equation takes the form

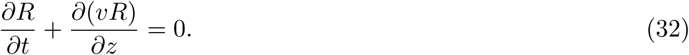

Here, lipid flows are of two origins: they can be generated by pressure gradients along the membrane tube or induced by dragging the filament with respect to the membrane. Consider a narrow segment of the membrane tube with radius *R*, width ∆*L* and surface area *S* = 2*πR*∆*L*. The energy of this tube segment is ∆*E* = *ϵ*(*R*)∆*L* where *ϵ*(*R*) is the energy per unit tube length. For the considered two-dimensional flow, the line pressure can be defined as

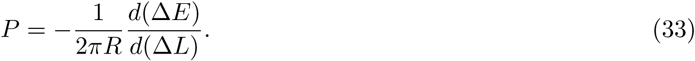

It specifies the force applied to a unit circumference element in the cross-section of the membrane tube. Because the surface area ∆*S* of the membrane is conserved, an increase of the segment width by *d*(∆*L*) must be compensated by a change in its radius by *dR* = −(*R*/∆*L*)*d*(∆*L*). Therefore, the elastic energy of this segment will be changed by

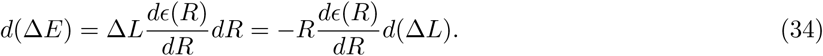

Hence, the pressure is

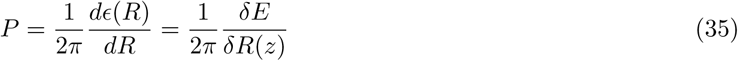

where the functional derivative of the total energy *E* taken at position *z* is introduced. A gradient of the pressure will induce a membrane flow with the velocity

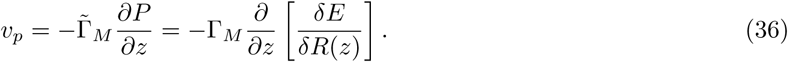

Here, 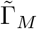 is a kinetic coefficient that specifies the effective mobility of the membrane and 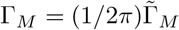.

If the membrane tube is covered by a moving filament, it will drag the tube because of the viscous friction between the filament and the membrane. We will determine the drag component *v*_drag_ of the flow velocity below using the polymer description for the filament. Hence, the evolution equation for the membrane in the continuum limit will be

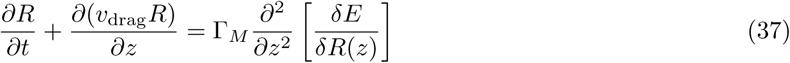

This evolution equation has to be complemented by a boundary condition at the ends of the membrane tube. Depending on the physical set-up, various conditions can be applied. In the present study, we will assume that the membrane tube is long and the filament occupies only a small part of it. At both ends, the tube has the equilibrium radius *R*_eq_.

For numerical simulations, membrane discretization should be additionally performed. We divide the tube into a sequence of disk segments *j* of radius *R*_*j*_, each of width ∆*L* = *d*_M_. Note that such disks are only introduced for the discretization means; they are immobile and lipid flows can enter or leave them. To determine *v*_drag_, suppose that within a disk *j* there are some beads *i* with axial velocities *u*_*i*_, whereas the flow velocity of the membrane is *v*_*j*_. Then, each bead experiences a viscous friction force *f*_*i*_ = (*v*_*j*_ − *u*_*i*_)/Γ from the membrane and, reciprocally, the opposite force −*f*_*i*_ acts on the membrane. Therefore, the total drag force acting on the tube segment is

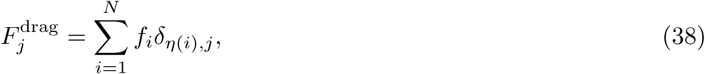

where the Kronecker delta symbol *δ*_*η*(*i*),*j*_ restricts the sum to the beads belonging to membrane disk *j* (see Eq. 24). On the other hand, the forces must be balanced for each bead *i* and therefore the viscous friction force *f*_*i*_ should be equal to the force applied to this bead, i.e.

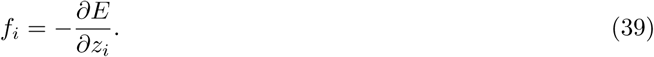

Moreover, the drag force applied per unit membrane area is 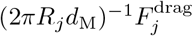 and, therefore, the velocity of the induced lipid flow is 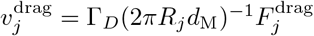 where Γ_*D*_ is a kinetic coefficient. Because this coefficient has the same dimensionality as Γ_*M*_, in numerical simulations we shall assume that Γ_*D*_ = Γ_*M*_.

Combining all terms,

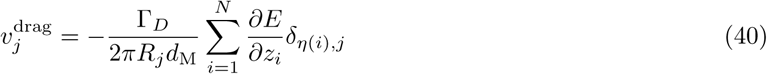

This expression along with Eq. 37 completes the description of membrane evolution.

The discretized version of the evolution equation (Eq. 37) is

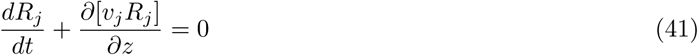

with

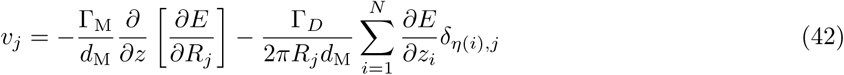

where the sum is taken over all ∆*N*_*j*_ beads *i* that are currently located within the disk segment *j*. The first term takes into account lipid flows induced by the pressure gradient and the second term accounts for the filament drag of the membrane. In the above two equations, a concise operator notation

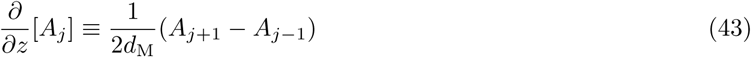

is used. Fixed radial boundary conditions are implemented by taking *R*_1_ = *R*_2_ = *R*_3_ = *R*_*N−*2_ = *R*_*N−*1_ = *R*_*N*_ = *R*_eq_.

#### 2.2.4 Characteristic time scales

The physical connection between the filament and the membrane is mediated through the PH domains, which are partially immersed into the membrane [14] and/or bound to single phosphoinositide lipids [13]. Since the viscosity of a lipid bilayer is typically 1000 times higher than that of water, the mobility of filament beads along the membrane, *i.e.* in the *φ* and *z* directions, should be controlled by their interactions with the membrane. Therefore, we assume that the mobility of the beads is approximately the same as that of the individual lipids. Diffusion constants of single lipids can sensitively depend on the physical state and the composition of a biomembrane. For our numerical estimates, we choose *D* = 1 nm^2^/µs, see [39]. The mobility of a lipid (or filament bead along the tube) is linked to its diffusion coefficient through the Einstein relationship and, therefore, Γ = *D*/k_B_T. This means that an applied force of *F* = 1 k_B_T/nm= 4.1 pN could generate a velocity *v* = Γ*F* = 1 nm/µs. Under the action of such a force, a bead would need only 10 µs to move over a distance of 10 nm.

As opposed to motion along the tube, radial bead motion is constrained to follow the local membrane radius, which is coupled to membrane flow by the continuity equation. The membrane mobility coefficient Γ_M_ specifies the local lipid flow velocity induced by application of a pressure gradient along the tube. This coefficient can depend only on the viscous properties of the lipid bilayer forming the membrane. The dimensionalities of Γ_M_ and Γ differ by a factor representing a square of acharacteristic length. Choosing it as the length *d*_lipid_ of a lipid, we obtain, by order of magnitude, 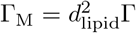. Using Eqs. 37 and 11, we find that

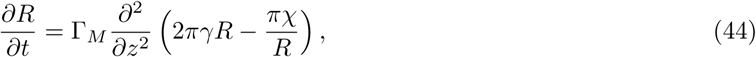

where terms with higher spatial derivatives have been dropped. For small deviations *δR* from the equilibrium tube radius *R*_eq_, this equation can be linearized to obtain

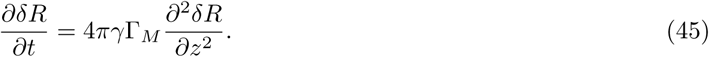

Thus, the characteristic time scale *t*_*L*_ for relaxation of membrane radius perturbations with the length scale *L* is

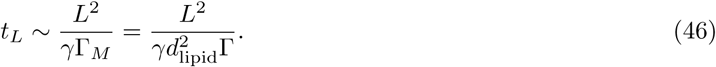

where the numerical prefactor has been dropped.

If the membrane tension is *γ* = 0.01 k_B_T/nm^2^ and *d*_lipid_ = 1 nm, perturbations with the size of *L* = 10 nm will relax within the characteristic time *t*_*L*_ = 10 ms. This membrane relaxation time increases, however, already to 1 s if the perturbation length *L* is 100 nm. Because the elasticity of the filament is not too different from the membrane, relaxation times for the tube covered by the filament should not be much different from these estimates. As we see, polymer bead motions are much faster in comparison to relaxation processes in the membrane. This is because the latter ones are nonlocal, i.e. accompanied by lipid flows involving the entire membrane tube.

It should be stressed that our estimates of characteristic times are very rough and should be used only as orientation. Differences by an order of magnitude are well possible. They may arise because diffusion coefficients of lipids can vary substantially, depending on the membrane chemical composition and its physical state. Moreover, strong simplifications have been made and we have also systematically omitted all numerical factors.

## 3 Methods

### 3.1 All-atom simulations of a tetramer

To estimate the flexibility of the stalk, an explicit-solvent simulation was performed for a stalk tetramer construct. The simulation was initialized from the tetramer crystal structure [11] and contained for each monomer residues 321-499 and 643-698 that comprise the four helices of the stalk. A 5 residue SSGSS linker was inserted between residues 499 and 643, where the PH domain is cut out. K^+^ and Cl^*−*1^ ions were used to neutralize the system and provide a salt concentration of 0.15 M along with the TIP3P water model. Periodic boundary conditions were employed with a box measuring 93×93×169 Å, which contained 15,708 protein atoms, 133,482 solvent atoms, and 252 ions. Simulations were performed with Acellera ACEMD [40] using the CHARMM36 forcefield [41]. Details of the simulation are as follows: NPT ensemble, temperature 310 K, Langevin thermostat, Berendsen barostat at 1 atm, holonomic restraints on hydrogen bonds, hydrogen mass scaled by factor of 4, timestep 4 fs, PME electrostatics, grid spacing 1 Å, cutoff 9 Å, switching at 7.5 Å. The simulation box was equilibrated for 10 ns under the NVT ensemble and then 10 ns with the NPT ensemble before a long production run of 3 µs.

### 3.2 Coarse-grained simulations of a filament

We used a well characterized structure-based model (SBM) (often termed a Gō-model) [42] to simulate a 17-dimer filament, sufficient linear in length to investigate twist angle fluctuations. The filament representation is single-bead-per-residue and residues close together in the native configuration (that is, part of the native contact map) are given attractive interactions. The contact map is determined by the Shadow criterion [43]. All other pairwise interactions are strictly repulsive. The native state (Fig. 4) was created by connecting 17 dimers together using identical interfaces 1 and 3. The coordinates of the dimer and of the tetrameric interface were taken from the crystal structure of the tetramer [11]. See Fig. 3A for the definition of the interfaces. Topology files compatible with GROMACS 4.5 [44] were created using the SMOG v2.0 software package [45] with the provided SBM calpha+gaussian template, which implements the 0.5 Å width Gaussian contact model of Lammert *et al.* [42]. The temperature in the simulation (1.05 reduced units, GROMACS temperature 126 K) was chosen such that the ensemble average root mean square deviations averaged over all C_*α*_ atoms matched between a tetramer simulated with the SBM and the explicit solvent model described above.

**Figure 4:**
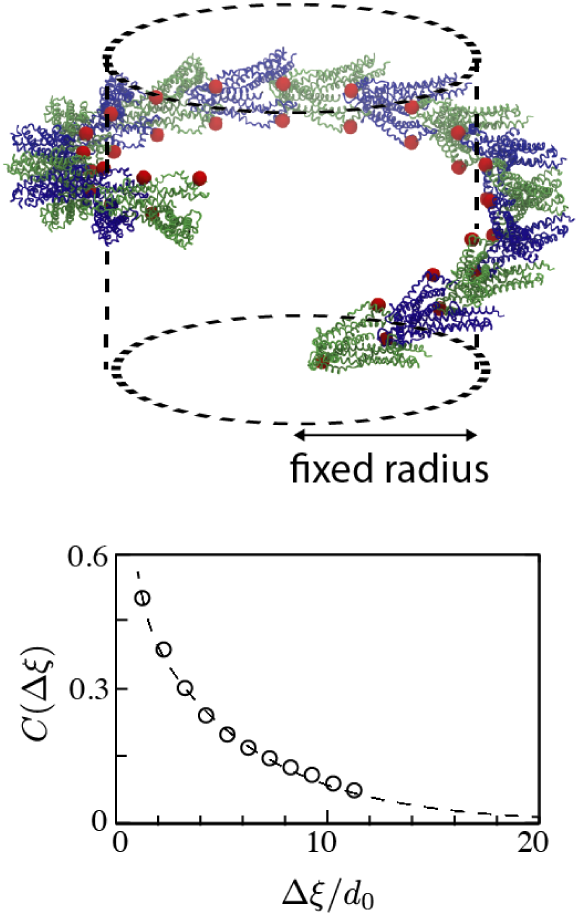
Estimating geodesic stiffness *β* for the stalk filament from coarse-grained molecular dynamics simulations. (Top) The simulated 17-dimer filament used to determine the twist correlation function *C*(∆*ξ*). Dimers are shown as alternating green and blue for clarity. The employed connection (53) between the correlation length and *β* holds for an oriented stalk filament on a rigid rod. These requirements are implemented by constraining the radii of all residues shown as beads to the fixed radius of 15 nm. (Bottom) The correlation function of twist fluctuations as a function of the filament arc length: simulation data (circles) and its single exponential fit (dotted line).

The twist angle correlation length along the filament follows Eq. 53 under two assumptions: 1) a small twist with *h≪ r* and 2) the filament is wrapped around a rigid tube. The native filament has *h* = 1.6 nm, and thermal fluctuations do not drive *h* greater than 3 nm, thus, the first assumption is valid. The validity of the second assumption is assured by strongly constraining the radius of the filament. One residue per monomer (Leu652) is constrained via the potential, 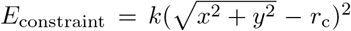, where k = 1000 k_B_T/nm^2^ and *r*_c_ = 15 nm. *r*_c_ is chosen to be consistent with the native filament’s radius (17 nm from the *z*-axis to a dimer’s center of mass). An additional consequence of the constraint is that it maintains the filament orientation (i.e. 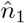 in Fig. 1). The twist angle correlation converged within 3× 10^8^ timesteps as there was no apparent difference between calculations using either the first half or the second half of a trajectory containing 6 × 10^8^ timesteps.

### 3.3 Mesoscopic polymer simulations

Simulations with the polymer model describe the coupled dynamics of an elastic filament attached to lipids on the outer leaflet of a membrane tube. Specific to dynamin are the input values of the filament elasticity (Table 1) and the geometry of the filament, i.e. beads are separated from each other by 5.6 nm and from the membrane by 8.5 nm.

**Table 1:**
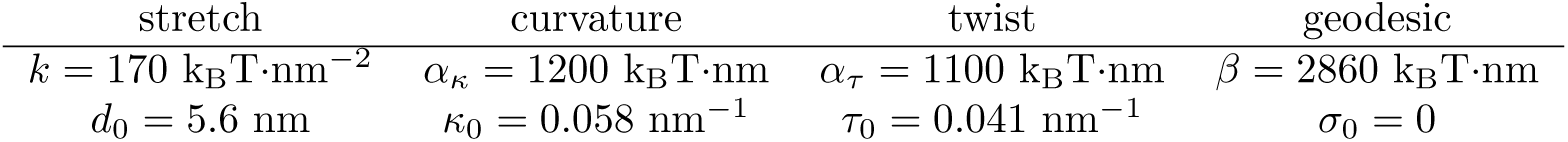
Elastic parameters of the dynamin filament

The equations of motion were implemented in an in-house version of the GROMACSv4.5.3 molecular dynamics package. The code is available upon request. All interaction potentials besides the harmonic nearest-neighbor interaction were implemented by adding code to handle the elastic forces. Internally, the units for energy were k_B_ × (310 K), length nm, and time µs. The energies are discussed in detail in Section 2.2, and the forces are worked out in the Supplementary Material. The elastic constants for the dynamin filament were set to those shown in Table 1. The membrane tension *γ* and stiffness *χ* took various values. The discretized membrane contained disks of width *d*_M_ = 4 nm, and the interaction distance between a membrane disk and the filament was *a* = 8.5 nm. The filament beads were separated by *d*_0_ = 5.6 nm. The interaction parameters were *d*_rep_ = 9 nm, *ϵ*_int_ = 20 k_B_T/nm^3^, and *ϵ*_rep_ = 1 k_B_T, but the dynamics described here is not sensitive to their specific value. For simplicity, the employed value of *ϵ*_int_ strongly constrains the deviation of the filament and membrane (|*R*(*z*) − *ρ*(*z*)| ≲ 1 Å. If attractive interactions between neighboring turns are included in a future model, choosing *ϵ*_int_ ~ 1 k_B_T/nm^3^ may be appropriate as it allows a few nanometers of variation consistent with the flexible attachment of the stalk filament to the PH domains. Simulations of long filaments contained *N* = 200 beads, where each bead corresponds to a dynamin dimer. With an average separation between beads of *d*_0_ = 5.6 nm, the total length of such a filament is 1.12 µm when measured along the filament, but, due to helical wrapping, only 100-300 nm when measured along the tube axis depending on the pitch.

The mobility for motion of filament beads over the membrane tube was Γ = 1 nm^2^/µs/k_B_T, i.e. similar to that of single lipids within the membrane. The mobility coefficient for the membrane was Γ_*M*_ = 1 nm^3^/µs/k_B_T. The boundary conditions for the membrane tube were that the radii of the first three and the last three discs were fixed to be the equilibrium radius *R*_eq_ of the bare membrane tube. The filament never reached the ends of the membrane.

For each system snaphot, the pitch was calculated by taking the average pitch of the central 21 beads (approximately a full turn). The local pitch for bead *i* is calculated using the approximate relationship

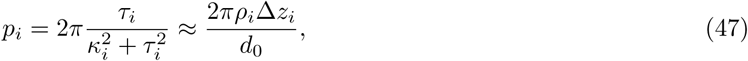

which is valid for ∆*z*_*i*_ *≪ d*_0_. The middle membrane tube radius for a given minimum energy shape was taken as *R*_min_ + *a*, where *R*_min_ is the narrowest membrane disk along the tube.

#### 3.3.1 Minimal-energy shapes

In an equilibrium state, lipid flows in the membrane are absent and such a state cannot be affected by them. The equilibrium shapes correspond to a minimum of the energy. Therefore, such calculations can be expedited by running simulations where the membrane tube obeys a simplified evolution equation

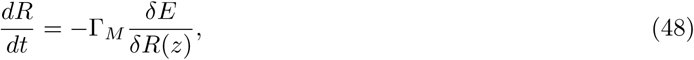

rather than Eq. (37). Note that the boundary conditions are still the same; the membrane tube has the equilibrium bare radius *R*_eq_ at its ends. The equation of motion for the filament remains unchanged.

For each simulation, the system was initialized as a filament with the pitch of 15 nm wrapped around a membrane tube with the equilibrium bare radius *R*_eq_. A simulation carried out for 1 × 10^8^ time steps at a low temperature (~0.05 k_B_T) and followed by 1 × 10^7^ time steps at *T* = 0 was sufficient to reach the minimum. This corresponds to roughly 15 minutes on a single core. Small thermal noise was introduced because, in the complete absence of thermal fluctuations, the system sometimes became trapped in local minima characterized by a membrane bulge and a large pitch bounded on either side by filament regions with a small pitch (similar to the barriers described in ref. [26]).

#### 3.3.2 Thermal fluctuations

Each simulation was initialized at a minimmal energy shape. The membrane flows were treated as described in Section 2.2.3 and the boundary membrane disks were held at a fixed radius *R*_eq_. Each simulation was run for a total of 1 × 10^9^ time steps where the time step was set to 0.01. This corresponds to roughly 1 day on a single core. The actual simulated duration can be estimated by using the mobility which was measured in µs. 1 × 10^9^ × 0.01 = 1 × 10^7^ µs or 10 seconds.

To determine mean-squared fluctuations, each snapshot yielded a single radius and pitch value for the averaging. The convergence was verified by checking that the averages and the mean-squared fluctuations agreed between the first half of the trajectory and the second half of it. The pitch was calculated as described above, but the radius determination was changed to be an average over the radii of the same central 21 beads used to calculate the pitch. Supplementary Movie 3 shows a 150 ms excerpt from simulations for three different membrane tensions.

Supplementary Movie 2 was run under the same conditions, except that the initial condition was a uniform tube of radius *R*_eq_ and only weak thermal fluctuations were introduced (*T* = 15 K).

## 4 Results

### 4.1 Estimation of elastic parameters from microscopic simulations

The high resolution structural information available for the stalk filament can be connected to a mesoscopic description of the filament as an elastic ribbon through molecular simulation. The nearly identical structures of the stalk tetramer inside a constricted stalk and of the isolated tetramer [8] suggest that constriction and relaxation of the filament radius is mediated by only subtle motions within the tetramer.

The fundamental repeating unit in the filament is the stalk dimer, which is stabilized by interface 2, while successive dimers connect together through two tetrameric interfaces 1 and 3 [11] (Fig. 3A). We expect that the dimer interface should be relatively rigid compared to the tetrameric interface for the following reasons: 1) Purified dynamin protein elutes as a dimer and higher oligomers but never monomers [9], 2) crystal structures of other related proteins show a consistent interface 2 [9, 46, 47], and 3) the first eigenmode of a principle component analysis of stalk tetramer simulations corresponds to bending/twisting about the tetrameric interface (Supplementary Fig. S2). Changing the angle *ϕ* defined in Fig. 3A from 15° to 25°, changes the membrane’s inner lumen from 11 nm to 2.5 nm (Supplementary Fig. S2). An additional advantage of this rigid dimer description is to provide a natural coarse-graining for the polymer model introduced in Section 2.2.

#### 4.1.1 All-atom MD simulations

In order to characterize the flexibility of the tetramer interface, we performed all-atom explicit-solvent molecular dynamics simulations of the stalk tetramer (see Methods for details). These simulations are similar to those in ref [9], but are initialized from the now available tetramer crystal structure [11].

Computational expense limits the calculations to a small piece of the filament (Fig. 3B), whereas the elastic degrees of freedom *κ* and *τ* are defined for a long continuous ribbon. Thus, a way of mapping the conformation of a tetramer onto *κ* and *τ* is needed. We do this by noting that a repeated, shape-conserving coordinate transformation Θ (see Supplementary Section 3 for details) always yields a helix. For a given tetrameric interface, Θ can be found by computing the transformation necessary to move a dimer within a tetramer from its position to the least root-mean-squared-deviation (RMSD) fit to its partner dimer.

Thus, a given tetramer configuration provides a transformation Θ, and repeatedly applying Θ generates a continuous helix with a particular radius *ρ* and pitch *h*. In turn, these *ρ* and *h* define the curvature *κ* and the twist *τ* corresponding to that snapshot by using Eq 4. For example, the transformation Θ moving one dimer onto the other in the tetramer crystal structure [11] creates the stalk filament shown in Fig. 1B. Further, computing the distance between the centers of mass of the dimers gives local changes in the stretch *d*.

The potential of mean force of the tetramer viewed along either *ρ* and *p* or *κ* and *τ* had a single dominant minimum (Fig. 3F). The elasticity of the stalk filament can be estimated from the distributions of stretching *d*, bending *κ*, and twisting *τ*, observed in the simulations (Fig. 3D/E). At thermal equilibrium, the fluctuations of local curvature and twist of a free filament (without the membrane) should obey Gaussian distributions (see Supplementary Section 4 for details)

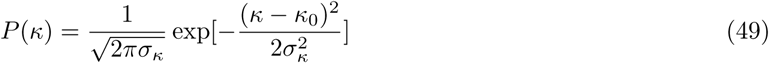

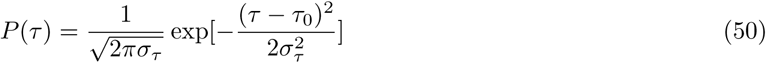

with the variances

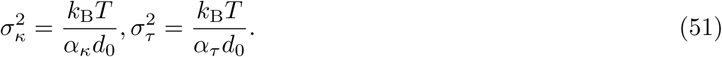

Thus, the peaks of the Gaussian fits should provide the spontaneous curvatures and the dispersions should give the elasticity moduli. Applying this to the MD simulation data, the numerical values shown in Table 1 were obtained.

The non-vanishing spontanous curvature *κ*_0_ means that dynamin has an intrinsic ability to bend the membrane upon which it is bound, whereas the condition *τ*_0_ *>* 0 implies a right-handed helix is formed. The small variation in the distances between centers of mass of the dimers *d* indicates that the filament is only weakly stretchable or compressible (Figure 3D). Note that clustering *d* with either RMSD, *κ*, or *τ* was not able to account for the small shoulder at *d >* 5.8 nm.

#### 4.1.2 Structure-based coarse-grained simulations

The remaining elastic coefficient, i.e. the geodesic curvature modulus *β*, could not be determined from the MD simulations for the tetramer. To find it, longer filament fragments were simulated in order to follow relative changes between tetramers. As shown in Supplementary Section 5, this elastic modulus can be determined by analyzing the correlation functions for fluctuations of the twist angle *θ* for a filament coiled over a rigid rod. The correlation function has the form

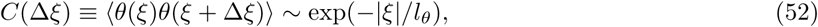

where the correlation length *l_θ_* for fluctuations of the twist angle about a stiff rod of radius *ρ* in the weak-twist approximation is

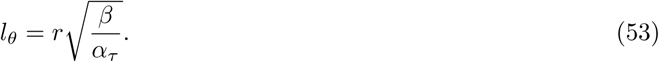

While a tetramer was sufficient to calculate the local twist, determining the correlation length of twist fluctuations *l*_*θ*_ requires a stalk filament longer than *l*_*θ*_.

In order to reach the necessary time scales to converge to the equilibrium distribution, we use a simplified C_*α*_ representation of a 17-mer filament and a structure-based model (SBM) [48, 45]. Structure-based models (also termed Gō-models) define a known accessible structure as a global energetic minimum and have been shown to capture the slow deformation modes defined by the network of inter-residue interactions [49].

The SBM for the filament was created using the tetramer crystal structure as the global minimum for each tetramer. Next, tetramers were connected together into a filament by repeatedly tiling the tetrameric interface, which created a helix with radius 17 nm, pitch 10 nm, and *h* = 1.6 nm [11] (Figure 4). Since there is no explicit membrane representation, the rigid rod assumption inherent in Eq. 52 and the perpendicular orientation assumed by the geodesic curvature were maintained by tightly, and identically, constraining the radii of the subset of atoms shown as spheres in Fig. 4 relative to the tube axis. For this helix, the weak twist approximation holds since *h ≪ ρ*.

By analyzing the computation data, the value *l*_*θ*_ = 5.0*d*_0_ was obtained through an exponential fit to the correlation function *C*(*τ* (*ξ*)*−τ* (*ξ* +∆*ξ*)) (Figure 4). This correlation length leads to an estimate of the geodesic curvature modulus *β* = 2.6*α_τ_* = 2860 k_B_T*•*nm.

#### 4.1.3 Comparison with elastic properties of other polymers and of membrane

It is instructive to compare the calculated elastic moduli for the stalk filament in Table 1 with the stiffness parameter of the of other biopolymers and the membrane. Ordinary materials have a Poisson ratio *ν* in the range 0 *< ν <* 1/2 [29], which implies 2*α_κ_*/3 *< α_τ_ < α_κ_*. This condition is satisfied by the calculated values of *α*_*κ*_ and *α*_*τ*_. Furthermore, note that the assumption that *α_κ_* = *α*_*τ*_ is reasonably good, although we shall not use it.

The persistence length *l*_*p*_ of the stalk filament is determined by the normal curvature elastic modulus as *l*_*p*_ = *α*_*κ*_/k_B_T= 1.2 µm. For comparison, the persistence lengths for actin and microtubules are 17 µm and 5000 µm, respectively. Therefore, we find that dynamin filaments are much softer than these other biopolymers. Such softness may be needed for the dynamin protein to act as a curvature sensor of the membrane. This is consistent with the presence of a soft mode in the tetramer (Supplementary Fig. S2).

An experimental measurement of the persistence length of long, dynamin-coated membrane tubes yielded a persistence length of 37 µm [50]. However, the above-defined correlation length refers to bending of free filaments along the filament coordinate *ξ*, whereas the measured persistence length corresponded to bending of dynamin-coated membrane tubes along their axial directions. The axial persistence length for the tubes may be strongly influenced by dimerization of GTPase-BSE domains that leads to cross-links established between the helical turns.

However, the dynamin filament should also constrict the membrane [18]. Therefore, the filament needs only be sufficiently stiff to remodel its membrane template, with any further stiffness potentially hindering the function. The relative stiffness of the stalk filament and the membrane can be compared by considering the dimensionless ratio *α*_*κ*_/*χp* [18]. For a pitch of *p* =10 nm and the membrane bending stiffness *χ* = 20 k_B_T a ratio of 6 is obtained, which is not much larger than one. This implies that the elasticities of both the filament and the membrane have to be taken into account.

It can be noted that the geodesic stiffness constant *β* is about 2.5 larger than the stiffness constants *α*_*κ*_ and *α*_*τ*_ for variations of normal curvature and twist. Therefore, even relatively small geodesic curvatures developing near free ends of a filament (cf. Fig. 7C below) can lead to substantial contributions to the deformation energy of the filament. The geodesic curvature effects must therefore be taken account when short dynamin filaments are examined.

### 4.2 Equilibrium shapes of filament and membrane

If the length of the filament is fixed, the equilibrium state should correspond to the minimum of the total energy *E* = *E*_F_ + *E*_M_ + *E*_int_. Thus, the equilibrium shape of the filament occupying a part of a deformable membrane tube is generally determined by the variational equations

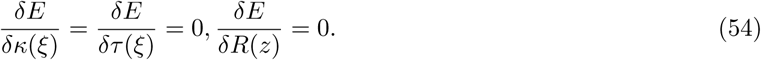

For a free membrane tube or free filament, the shapes can be easily determined with Eq. 54. We find that a free filament takes the values of its spontaneous curvature *κ*_0_ and spontaneous twist *τ*_0_, giving a radius of 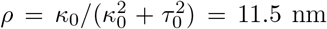 and a pitch of 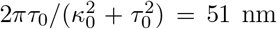. If such a filament were covering a membrane tube, its radius would have been *R* = *ρ − a* = 3 nm. A free membrane tube takes an equilibrium radius *R*_eq_ that balances the bending stiffness and tension (Eq. 12). As a point of reference, the membrane stiffness *χ* varied in the experiments between about 12 and 50 k_B_T and membrane tension *γ* between 0.005 to 0.025 k_B_T/nm^2^ [21]. For example, if *γ* = 0.01 k_B_T/nm^2^ and *χ* = 20 k_B_T, we have *R*_eq_ = 32 nm.

When interactions are present, explicit equations (Eq. 54) can be derived and used to compute the equilibrium shapes of the filament and the membrane. However, they turn out to be too complicated for closed form solutions. Instead, detailed equilibrium shapes will be determined in Section 4.4.2 by direct numerical integration of the evolution equations for the polymer model. Here, we will first perform a simplified analysis that yields only the radii of the filament and the membrane far from the ends.

#### 4.2.1 An approximate analysis for long filaments

Suppose that a filament of length *L*_f_ occupies a part of a long membrane tube with a total length *L* (Fig. 5A). It should be expected that the shapes of the filament and of the membrane are uniform outside of an interface region with the length of the order of *R*_eq_ near the filament end. If we neglect the interface contributions to the total energy, we have

**Figure 5:**
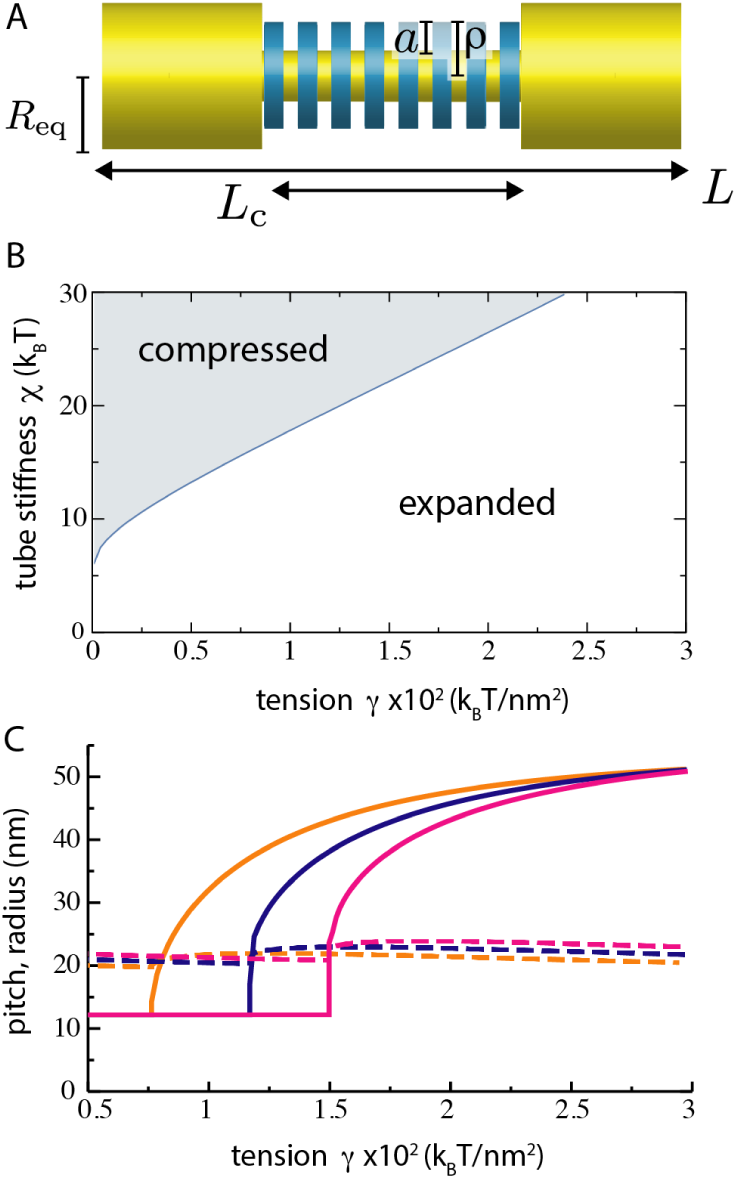
Tension controlled filament compression. A. Variables and parameters used in the approximate treatment. B. Boundary of filament compression. Inside the shaded region, the filament has the minimal possible pitch *p*_*min*_. C. Dependences of the equilibrium pitch *p* (solid) and radius *ρ* (dashed) of the filament on membrane tension *γ* for *χ/k*_*B*_*T* = 16 (orange), 20 (blue), and 24 (magenta). Other parameter values are as given in Table 1.

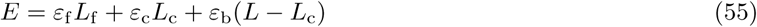

where *L*_c_ is the length of the membrane tube which is covered by the filament and *ε*_f_, *ε*_c_, *ε*_b_ are the elastic energies per unit length for the filament, the covered membrane, and the bare membrane, respectively. The interaction energy vanishes, *E*_int_ = 0, if, in the filament-covered part of the tube, *R* = *ρ* − *a*. The bare membrane tube has the energy per unit length of

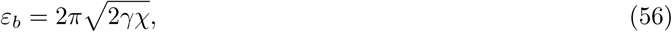

which is obtained by substituting *ρ* = *R*_eq_ into Eq. 11. The energy densities *ε*_f_ and *ε*_c_ are given by Eqs. 7 and 11. According to Eq. 14, the total axial length of the membrane tube covered by a filament of arc length *L*_f_ with radius *ρ* and pitch *h* is

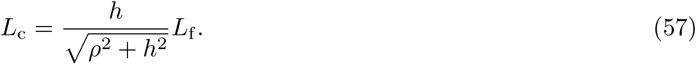

Summing up all contributions, we find that the total energy of the system is

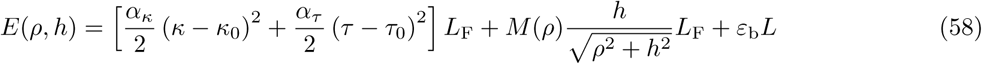

where

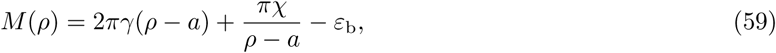

and recall that *κ* and *τ* can be expressed in terms of *ρ* and *h* in (Eq. 4).

The first term in Eq. 58 is the elastic energy penalty for deforming the filament to a curvature and a twist that are different from their spontaneous values. The second term is the penalty for compressing the membrane tube to the radius smaller than its equilibrium radius *R*_eq_ over the filament-covered part. The equilibrium values of *ρ* and *h* satisfy a system of variational equations

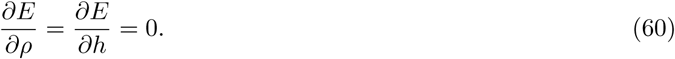

The energy per a dynamin dimer of length *d*_0_,

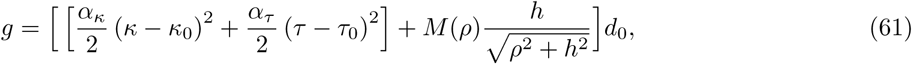

is plotted as a function of *ρ* and *p* = 2*πh* for different membrane tensions *γ* in Supplementary Fig. S3. For *ρ* = 20 nm and *p* = 12 nm at *γ* = 0.01 k_B_T/nm^2^and *χ* = 24 k_B_T, we have *g* = 5.9 k_B_T/dimer. Note that this is in the range of polymerization energy of dynamin, previously estimated to be 4-5 k_B_T/dimer [18].

### 4.3 Membrane-induced compression of the filament

Inspection of the second term in Eq. 58 reveals an important feature of the system: while keeping the filament radius constant, the energy can be lowered if the pitch of the filament is decreased. Since the filament is then more densely coiled, this would reduce *L*_c_, the length of the membrane tube over which it is spread. Effectively, the membrane generates a force that tends to compress the filament along its axis, which is counterbalanced by the filament’s twist stiffness. Formally, given a sufficiently stiff membrane, the pitch *p* = 2*πh* will vanish. In reality, the filament can be only compressed down to some minimal pitch *p*_min_ ~ 12 nm because of the finite size of the filament. Thus, the compression is manifested by a transition to such a maximally compressed filament state.

The boundary of the compression transition in the membrane parameter plane (*γ, χ*) can be found by setting *h* = *h*_min_ = *p*_*min*_/2*π* in the above variational equations (Eq. 60) and solving them. Fig. 5B shows this boundary when using the values in Table 1 as the elastic parameters of the filament. This line separates the parameter regions with maximal compression of the filament (*h* = *h*_min_) and with an expanded filament (*h > h*_min_). Interestingly, this structural transition in dynamin is taking place within the range of parameters characteristic for cellular biomembranes.

Moreover, by solving the variational equations, the pitch and the radius of the equilibrium filament can be determined for arbitrary membrane parameters (Fig. 5C). At large membrane tensions *γ*, the compression is weak and the filament pitch approaches its free filament value of 51 nm. As the membrane tension is decreased, this leads to compression of the pitch until *p*_min_ is reached.

It may appear paradoxical that less tense membranes exhibit stronger filament compression effects. To better understand this behavior, note that *M* (*ρ*) in Eq. 58 can also be expressed as

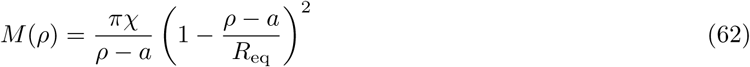

and, since *R*_eq_ *> ρ − a* and *R*_eq_ decreases as *γ* grows, increasing *γ* lowers the membrane energy. In other words, a tenser membrane has a smaller equilibrium radius, and thus requires less deformation to constrict it as the filament coils around it.

Whereas the pitch can dramatically change with variation in membrane parameters, the radius of the filament is remarkably constant. It stays close to *ρ* = 20 nm, which is smaller than the equilibrium radius *R*_eq_ of the free membrane tube, but larger than the equilibrium radius *ρ* =11.5 nm of the free filament. This can be compared with the filament radius of 18 nm in the “non-constricted” cryo-EM structure [38]. Additionally, optical-trap tube-pulling experiments have indicated that there is at most a weak dependence of the filament radius on the membrane tension [18]. We find similar behavior in polymer simulations where the full shapes of the filament and the membrane tube can be obtained.

### 4.4 Polymer model simulation results

In contrast to the analytical treatment of the previous section, the polymer model additionally accounts for the interface region at the filament ends. Thus, a first question is how long must a filament be to reach its asymptotic uniform shape? Determination of the minimum membrane radius as a function of filament length shows that the interior of filaments with *N >* 120 have reached their asymptotic shape (Supplementary Fig. S4). Therefore, in the following sections we describe the shape of “long” filaments with *N* = 200 (1.12 µm in linear length) wrapped around a membrane tube 1 µm in length. The membrane tube has boundary conditions that fix its ends to *R*_eq_ and the filament does not reach the ends.

#### 4.4.1 Filament-induced membrane flow

An illustration of the dynamics of filament-induced membrane flow and the corresponding membrane-induced filament compression is presented in Fig. 6 and Supplementary Movie 2. Here, the system is initialized in a non-equilibrium configuration where a compressed filament is wrapped around a uniform membrane tube of radius *R*_eq_ with *γ* = 0.01 k_B_T/nm^2^ and *χ* = 24 k_B_T.

**Figure 6:**
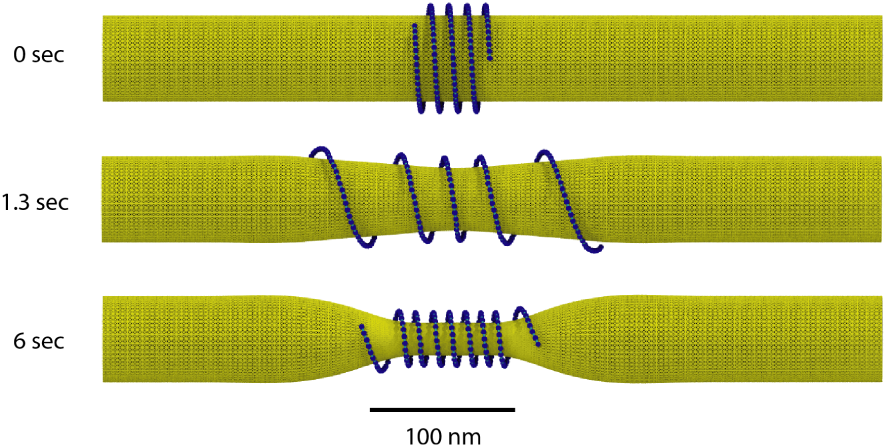
Filament relaxation toward its equilibrium shape accompanied by flows in the membrane. The initial condition and two snapshots are shown. The membrane was initialized with a uniform radius of *R*_eq_ and the membrane parameters were *γ* = 0.01 k_B_T/nm^2^ and *χ* = 24 k_B_T. The filament was initialized with a 12 nm pitch. Small thermal noise corresponding to 15 K is included to avoid local energy traps. Note that only 500 nm of the 1 µm membrane tube is shown. The full movie is available as Supplemental Movie 2.

As can be seen, the filament first quickly expands over the tube thus relaxing its pitch. Then, a more slow evolution process of tube constriction, accompanied by lipid flows in the membrane, sets in. Because the constricted membrane tube generates pressure compressing the filament in the axial direction, the pitch of the filament gradually decreases. Finally, the equilibrium state of the filament and the membrane is reached. The compression dynamics relaxes a 250 nm perturbation over ≈ 6 seconds, which is in agreement with the order of magnitude estimation of 6 seconds suggested by Eq. 46.

Remarkably, it can be noticed in the Supplementary Movie 2 that, when the filament contracts towards its final equilibrium shape, a bulge develops in the middle of the membrane. The interpretation of this effect is that the contracting polymer drags the lipids towards the middle of the tube. This induces lipid flows and, because of the mass conservation, a local inflation of the tube occurs. When the filament contraction is completed, such drag forces vanish and, through lipids redistribution by the flows, the membrane approaches its final equilibrium shape, being most strongly constricted in the middle.

#### 4.4.2 Full equilibrium shapes of the filament and membrane

The minimum energy shape of the filament on a membrane of stiffness *χ* = 24 k_B_T and tension *γ* = 0.01 k_B_T/nm^2^ is shown in Fig. 7A (see Methods for simulation details). The radius and pitch are constant within the filament except at the end, where the filament winds around a tube of growing radius. The dependencies of the filament radius *ρ* and pitch *p* on the coordinate *z* along the tube, together with the dependencies of the local normal curvature *κ*, twist *τ*, and geodesic curvature *σ*, are shown in Fig. 7B/C. As *ρ* is forced to follow the growing tube radius, the other quantities vary as well. The pitch of the filament is constant in its central part, but increases almost four-fold at its end. The geodesic curvature *σ* is non-vanishing in the interface region near the filament end.

**Figure 7:**
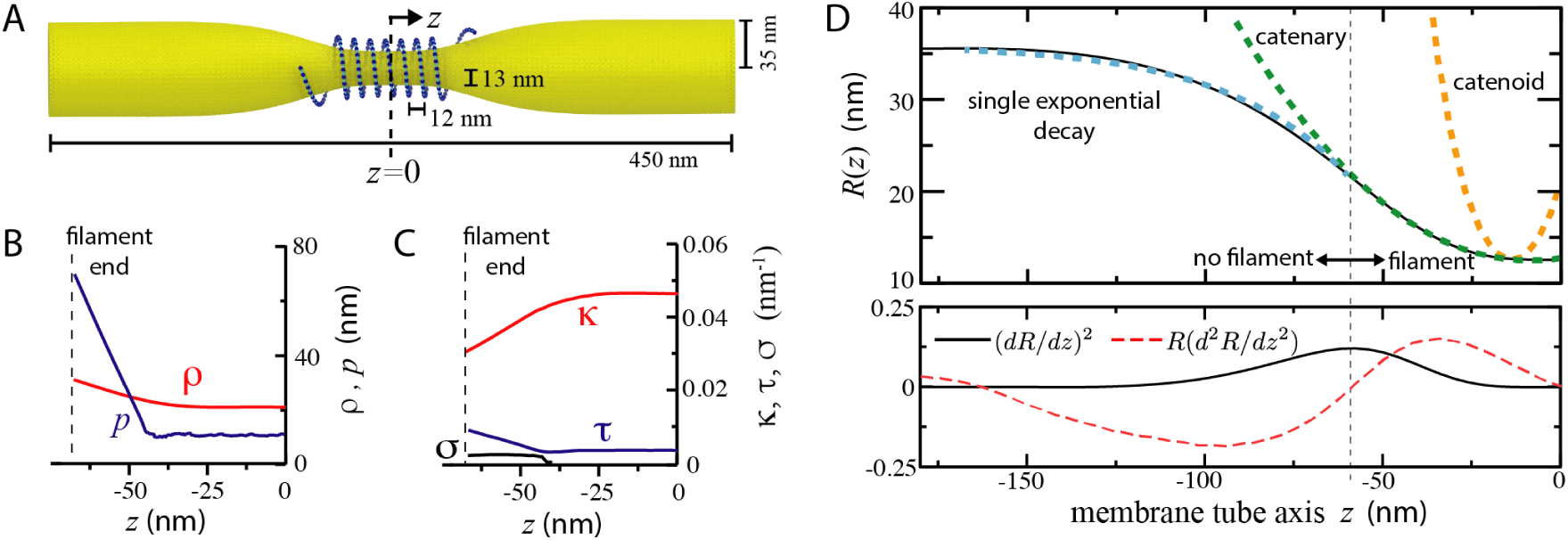
Full shape of the membrane/filament system. A) Example of an energy minimized configuration for a condensed filament with membrane parameters *χ* = 24 k_B_T, *γ* = 0.01 k_B_T/nm^2^. B/C) The corresponding dependencies of filament variables on distance *z* along the tube, including the interface region. D) The corresponding dependencies of the membrane radius and of its two derivatives along the tube in the interface region. Green dotted line is a catenary fit for the region with *d*^2^*R/dz*^2^ *>* 0 (Eq. 63). Cyan dotted line is the exponential decay function (Eq. 64). (D, lower panel) The value of (*dR/dz*)^2^, the relevant the small parameter in the locally-cylindrical helix approximation, is plotted for the membrane profile shown above. As discussed in the Supplementary Information, for a filament with its normal (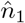) everywhere coinciding with the membrane surface normal, another quantity *R*(*d*^2^*R/dz*^2^)^2^ should also be small along the filament.

The corresponding profile of the membrane tube is analyzed in Fig. 7D. Far from the filament, the membrane tube takes a constant radius of *R*_eq_, and well within the filament-wrapped part, the membrane tube takes on a constant, compressed radius *R*_min_. For the chosen membrane parameters, the interface region within which the radius decreases from *R*_eq_ to *R*_min_ is approximately 150 nm long. Remarkably, the inflection point *z* = *z*_infl_, where *d*^2^*R/dz*^2^ changes its sign, is found to coincide with the end of the filament.

The profile of the filament-coated membrane part at the free end of the filaments is well approximated by a catenary curve for *z*_infl_ *< z < z_c_* (see Fig. 7D):

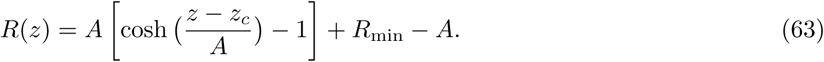

The two fitting parameters are *A*, the width of the catenary, and *z*_*c*_, the position along the axis where the radius starts to grow away from *R*_min_. For the catenary curve shown in Fig. 7, *R*_min_ = 12.2 nm, *A* = 126 nm, and *z*_*c*_ = −10 nm.

It should be stressed that the catenary fitting (Eq. 63) is different from a catenoid. The catenoid shape would have corresponded to the dependence (Eq. 63) with *A* = *R*_min_. This dependence is also shown, for comparison, in Fig. 7D. In contrast to a catenoid, an axially-symmetric surface defined by equation (Eq. 63) does not have a vanishing mean curvature.

In the bare part of the membrane tube (for *z < z*_0_), its shape is well approximated by an exponential dependence

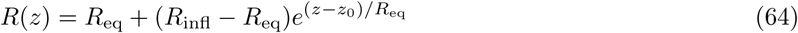

with the decay length of *R*_eq_, where *R*_infl_ is the tube radius at the inflection point *z* = *z*_0_.

With the membrane and filament profile in hand, we can check the condition (*dR/dz*)^2^ ≪ 1 that ensures the validity of the locally-cylindrical approximation for the membrane. As revealed by Fig. 7E for *γ* = 0.01 k_B_T/nm^2^, (*dR/dz*)^2^ is at most about 0.1 over the entire membrane tube. Thus, the locally-cylindrical approximation remains reasonably well applicable for the membrane even at the filament ends. The approximation worsens at the ends if the tension is further decreased. Because the filament radius is *ρ* = *R* + *a*, the condition (*dρ/dz*)^2^ ≪ 1, ensuring the validity of the locally-cylindrical approximation for the filament, is thus also verified.

#### 4.4.3 Filament compression by the membrane

The transition from the maximally compressed to the expanded filament, controlled by the membrane tension, has been found above through an approximate analysis of the continuum model in Section 4.2. Numerical simulations show that it is also present in the polymer description. Fig. 8 summarizes the results of numerical investigations of the filament compression effect.

**Figure 8:**
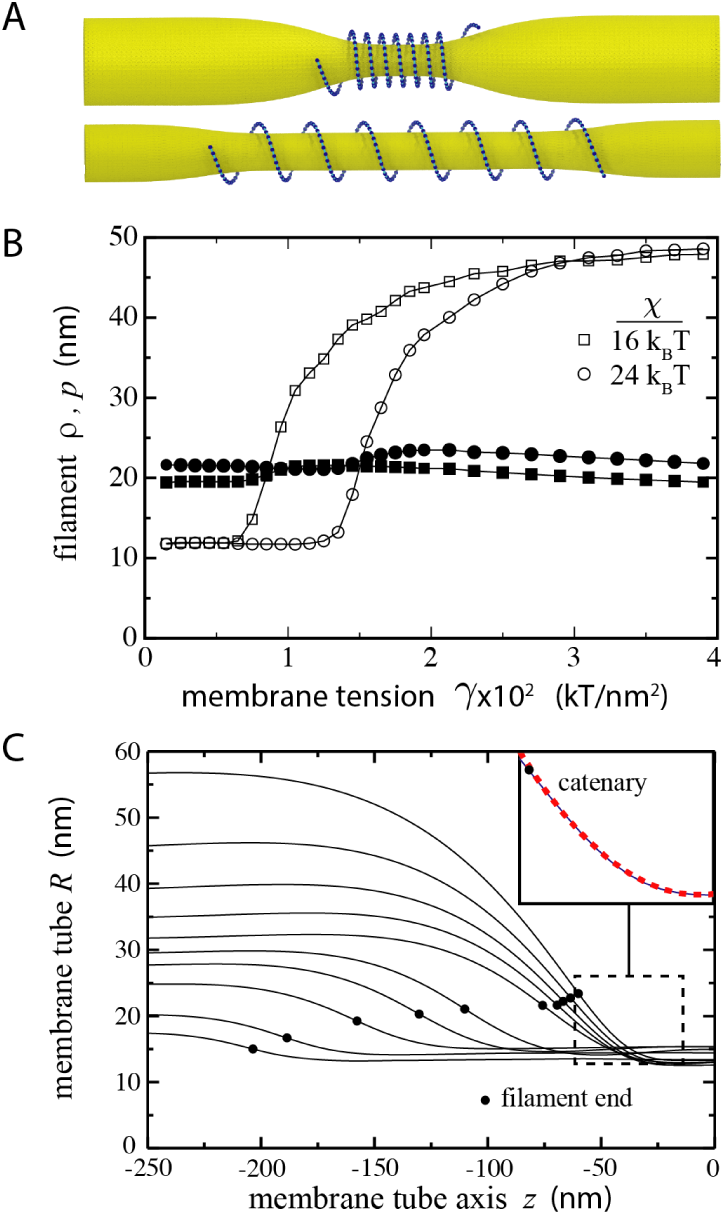
Membrane parameters influence the filament pitch. A) Two examples of energy-minimized configurations showing a condensed filament at low tension (top, *γ* = 0.01 k_B_T/nm^2^) and an expanded filament at high tension (bottom, *γ* = 0.025 k_B_T/nm^2^) for *χ* = 24 k_B_T. B) Dependencies of the radius (open symbols) and the pitch (closed symbols) of the filament on membrane tension *γ* for *χ* = 16 k_B_T (squares) and *χ* = 24 k_B_T (circles). The displayed values were obtained by averaging the pitch/radius over 20 beads in the middle of the filament; each symbol corresponds to one simulation. C) Equilibrium membrane tube profiles at *χ* = 24 k_B_T for different membrane tensions. From top to bottom: *γ* = 0.4, 0.6, 0.8, 1.0, 1.2, 1.4, 1.6, 2.0, 3.0, 4.0 × 10^*−*2^ k_B_T/nm^2^. The filled circles mark membrane contact points of the first bead of the filament. Since the profiles are symmetric, only the data for a half of the tube is displayed. The inset shows a catenary fit near the filament end for the lowest tension.

Below a characteristic critical tension *γ*_c_, depending on the membrane stiffness *χ*, the filament is compressed in its interior to the minimum pitch *p*_min_. For higher membrane tensions, the pitch grows with *γ* asymptotically approaching the maximum pitch of the free filament of 51 nm. Fig. 8B shows the dependencies of the equilibrium pitch and radius in the middle of the filament on the membrane tension. They are similar to those found in the continuum description (Fig. 5C). Note however that the polymer description does not yield a sharp transition at *γ* = *γ*_c_ since the interface region is now explicitly taken into account.

In contrast to the pitch, the radius is only weakly dependent on the membrane parameters. The combination of two effects can be here involved. First, the tube radius should tend to decrease with the growing tension, as already seen in the tension dependence (Eq. 12) of the equilibrium bare membrane tube. On the other hand, the decreasing coiling density, i.e. an increased pitch, may favor inflation of the tube and thus compensate the first effect.

Though the radius of the tube in the middle of the filament remains constant, the profile and the position of the interface region depend on membrane tension *γ*. In the bare part of the tube, the membrane radius approaches *R*_eq_, which decreases with increasing *γ*. Through the local interaction (Eq. 24) between the filament and the membrane, each bead feels a force acting along the tube direction and proportional to the gradient of the tube radius *dR/dz*. Such additional forces, acting on the beads at filament ends where the gradients exist, are responsible for filament compression by the membrane. Since the magnitude of the gradient decreases along with *R*_eq_ (Fig. 8C), the membrane-induced pressure eventually cannot counterbalance the intrinsic tendency of the filament to take on its spontaneous pitch, leading to filament expansion for *γ > γ_c_*. Notice that, even when *γ < γ_c_*, the end of the filament has larger pitch (Fig. 7B).

These results rely on the values of the elastic parameters obtained for dynamin in our MD simulations in Section 4.1. One may ask therefore whether such findings are strongly sensitive to the specific parameter values employed. To check this, minimum energy shapes for the filaments were additionally numerically determined by varying the elastic parameters, i.e. under the transformations *α*_*κ*_, *α*_*τ*_, *β → ωα*_*κ*_, *ωα*_*τ*_, *ωβ* (Supplementary Fig. S6). For a more stiff filament (*ω* = 10), we have found that the filament radius is approximately 15 nm and the pitch is larger than 30 nm across the whole range of membrane tensions. For a more pliant filament (*ω* = 0.1), a pitch compression similar to that at *ω* = 1 is found, whereas the radius increases above 30 nm at low membrane tensions. Experimental observations of dynamin-covered, low-tension, membrane tubes yield a filament radius and pitch near 20 nm and 10 nm, respectively [38, 5, 51, 52, 53]. Thus, the predicted equilibrium shapes of significantly stiffer or more pliant filaments appear to be inconsistent with the experimental measurements: the pitch would have been too large for stiffer filaments and the radius would have been too large for more pliant tubes.

#### 4.4.4 Thermal fluctuations

In the paper thus far, we have stayed in the deterministic limit or close to it. At nanometer scales, the effects of substantial thermal noise must also considered since they set the range of expected fluctuations and can shift the average values of observables away from their deterministic values. With the polymer model, thermal fluctuations of the filament are naturally included (Section 2.2.2). Explicit fluctuations in the membrane tube are still neglected by us, but we can notice that fluctuations in the beads will lead to fluctuations in the membrane because of the coupling of the filament to the membrane tube.

The effects of thermal fluctuations in numerical simulations of the polymer model are presented in Fig. 9. Here, a filament with 200 beads has been simulated sufficiently long to obtain the statistical averages. The entire simulation data was used to determine the mean radius 〈*ρ*〉 and the mean pitch 〈*p*〉 (see Methods for details). To determine the fluctuation intensities, local averaging of the radius and the pitch over 21 beads (note that a helical turn of radius 19 nm and pitch 12 nm contains 21 beads) was performed. Then, mean-square deviations of the smoothened variables were computed. See Supplementary Fig. S7and Supplementary Movie 3 for excerpts of trajectories with thermal fluctuations.

**Figure 9:**
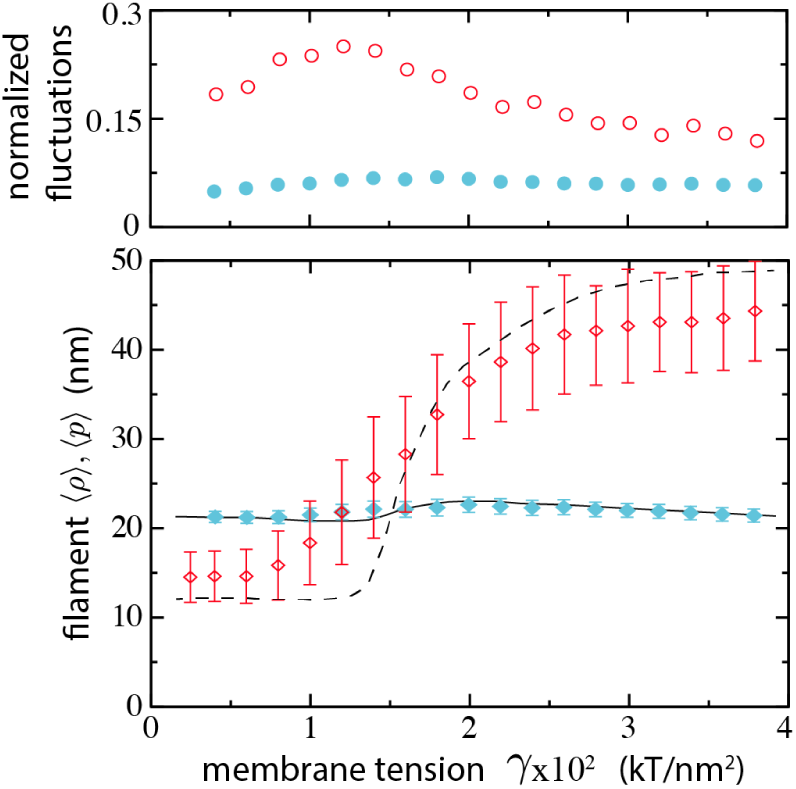
Thermal fluctuations in the polymer model. Data shown is for membrane stiffness *χ* = 24 k_B_T, *T* = 310K, and a filament containing 200 dimers. (Lower panel) Ensemble-average filament radius 〈*ρ*〉 is plotted as filled blue diamonds; average pitch 〈*p*〉 is plotted as open red diamonds. Solid/dotted line is the radius/pitch for the minimal energy shapes, i.e. zero temperature (see Fig. 8B). The bars indicate the root-mean-square deviations (RMSD) 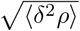 and 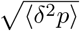 of the radius and the pitch from their mean values 〈*ρ*〉 and 〈*p*〉. (Upper panel) Relative fluctuations of the radius and the pitch. They are defined as the RMSD normalized by the ensemble averages. Filled blue circles show *σ*_*ρ*_/〈*ρ*〉; open red circles show *σ*_*p*_/〈*p*〉.

One can see that thermal noise has only a small effect on the radius of the membrane tube. Radial fluctuations are weak and, moreover, the statistical mean values of the radius are close to the deterministic predictions. The situation is, however, different for the pitch. Normalized fluctuations in the pitch are of about 20 percent in magnitude and have a maximum near the compression transition point. Moreover, the mean pitch differs substantially from that determined by energy minimization. This behavior can be understood by examining the energy landscape given by Eq. 58 (Fig. S3). The minimum determining the equilibrium shape lies inside an energy basin which is narrow along the *ρ* direction, explaining why the radius is well-defined and its fluctuations are weak. However, the basin is wide along the *p* direction. Therefore, fluctuations in the pitch are strong and substantial deviations between the mean pitch and the deterministic prediction are observed.

## 5 Discussion and Conclusions

We have developed a mesoscopic description for dynamin filaments coiled around membrane tubes. In our model, the filament is treated as an elastic polymer that naturally tends toward a helical shape. The elastic parameters of the filament could be determined from direct microscopic simulations of short fragments of it. Moreover, the model includes a deformable membrane tube with elastic energy given by the classical Helfrich description. The dynamical equations for membrane evolution, taking into account the membrane area conservation and lipid flows, are formulated as well. Thermal fluctuations at mesoscales are also resolved. Therefore, significant progress has been made in modeling the coupled behavior between dynamin and membranes.

Several modeling assumptions have been made. We considered membrane tubes with axially symmetric shapes and approximated by a cylinder with the slowly varying radius. The filament was assumed to have a locally-cylindrical helix shape. Because the radius of the filament follows that of the membrane, the latter condition is automatically satisfied when the membranes have the locally-cylindrical form. We have checked that, for the parameters characteristic for dynamin and biomembranes, the locally-cylindrical approximation can be justified for the entire filament, including also the parts near its free ends. Furthermore, we have found that the dynamin-coated part of membrane tube has a catenary shape at the filament end.

Another assumption was made concerning the interactions between the filament and the membrane. Physically, the filament consists of stalk domains connected by linkers to PH domains attached onto the membrane. Our assumption is that the linkers are flexible. Therefore, while keeping the filament near to the membrane, they do not locally orient it. Hence, the PH domains may be always orthogonal to the membrane, but their orientation is decoupled from that of the stalk domains. This assumption is supported by the fact that the linkers with the length of about 20 amino acids cannot be resolved in X-ray crystal structures. Moreover, the highest resolution cryo-EM map indicates random loop structures for them [8].

Furthermore, polymerization effects were not considered by us and the size of the filament was assumed to be fixed. Therefore, our results are formally applicable only close to the dynamin nucleation point, i.e. when the processes of filament growth or contraction are relatively slow. Thus, the determined equilibrium shapes may actually still adiabatically vary when the size of the filament gradually changes.

Our principal conclusion is that the elasticities of the filament and the membrane are of comparable magnitudes and therefore both of them must be considered when modeling is performed. Remarkably, our analysis has revealed that, within a realistic parameter range, the radius of the filament is not significantly sensitive to elastic properties of the membrane. Alone, this observation may seem to support a conjecture that the membrane is relatively soft. However, we have also found that the pitch of the dynamin filament is sensitively dependent on the membrane parameters. Generally, the membrane tends to compress the helical filament along its symmetry direction and the elastic membrane properties are crucial for this effect. In agreement with experimental data, we have found that, even in absence of GTP and of the motor operation, the elastic properties of the stalk filament are enough to constrict the membrane tube, but this constriction is not sufficient for membrane fission. In turn, the membrane acts on the filament and compresses it, tending to minimize the fraction of the tube length covered by dynamin.

In the future, the polymer model can be extended to consider further aspects of dynamin function, such as the motor operation powered by GTP hydrolysis. This will allow a study of the dynamin helix under the conditions that go beyond relaxation processes and fluctuations about the equilibrium states. Polymerization and depolymerization processes can also be included into the model. Hemifusion and scission are processes that go beyond an elastic description of the membrane, but coupling the polymeric filament with particle-based membrane models [54] would provide the possibility to follow the entire membrane scission process too.

During the scission process the helical dynamin oligomer is believed [3] to constrict its underlying membrane tube by creating forces that induce the constriction of the helix itself. The structural backbone of the dynamin oligomer consists of the stalk filament, formed by repeated connection of dimers through the tetrameric interface. Consistent with previous work [9], we have found that the crystallographic tetramer exhibits a soft mode and that the presence of such a mode effectively accounts for an adaptable radius of the stalk filament. The previously mapped energy profile appeared bimodal, with local energy minima at low pitch/low radius and high pitch/high radius. Possible reasons for the discrepancy are the use of a modelled tetramer or that the length of the simulations was only one tenth of the sampling here.

For a free helical filament in the absence of membrane and inter-helical interactions, the radius and the pitch are determined by the spontaneous curvature and spontaneous twist. As shown in Section 4.2, the radius of the free filament is 11.5 nm and its pitch is 51 nm. The equilibrium shapes of filaments on membrane tubes were then determined by using a continuum model, including elastic deformation effects for the membrane and the filament, and also the discrete polymer model for dynamin. We found that, when the filament is coiled around a tube, its radius is increased from 11.5 nm to about 22 nm, reflecting a contribution from the elasticity of the membrane. Hence, membrane tubes will be constricted to an inner lumen radius of 8-10 nanometers over a range of physiological membrane stiffnesses and tensions.

This range of tube diameters agrees with the determinations from various experimental techniques, including apo cryo-EM structure [38], negative stain EM [5, 51], in vitro fluorescence [52], and in vivo imaging of elongated necks in arrested clathrin-mediated endocytosis [53]. Similar tube diameters are even observed for a construct lacking the PH domain [52]. The correspondence between these experimental measurements and the presented model, which only includes the elasticity of the stalk filament, strongly suggests that the stalk filament alone provides the GTP-independent membrane shaping activity of dynamin.

The presented model shows a dependence of the filament pitch on the tension of the membrane and, remarkably, the strongest effects are exhibited by almost tensionless membranes. This is because the equilibrium radius of a bare membrane tube is large in the tensionless limit, and hence the relative constriction is stronger for such membranes. According to our results, the pitch varies from about 50 nm for high-tension membranes to 12 nm for low-tension membranes. Below the crossover tension, the filament gets maximally compressed and only subject to the excluded volume of the stalk, which was set to be 12 nm. The compression of the helical turns is reminiscent of the membrane-elasticity-driven clustering predicted for anisotropically-curved proteins [55].

In agreement with these results, most structural studies find a helical pitch under 15 nm for typical (in vitro almost tensionless) membrane tubes. Larger pitch values of up to 50 nm were not, however, so far experimentally observed, although they have been previously found in molecular dynamics simulations [9]. Even on stiff tubes, dynamin has been reported to take on pitches less than 25 nm [56, 57, 17]. This difference can be due to two additional effects which are not taken into account in the present model. First, it should be noted that polymerization processes [26, 28] are not yet included into it and therefore our results correspond to the filaments of fixed intrinsic length. The filaments may however tend to grow when such a process takes place. When the growth gets arrested at the end of a tube, the polymerization force [18] should appear leading to additional compression of the filament and thus to a shorter pitch. Second, GTP-induced cross dimerization of G-domains [56, 57, 17] should lead to additional strong interactions between helical turns, further favouring the compression to a shorter pitch. In the case of stiff, preformed membrane tubes, dynamin has been observed to display a distribution of larger pitches [56] when GDP-bound or with a mixture of apo and GDP-bound. Expansion pressure from the filament twist in the absence of strong G-domain dimerization seems to be a plausible explanation for this shape change.

Because the effects of both filament and membrane elasticity on the equilibrium shapes have been accounted for in the present study, previous modeling assumptions can be analyzed. As we have found, taking the tube radius as constant over a large range of membrane tensions [18] is a good approximation indeed. On the other hand, the tube neck shape near the ends of the filament likely cannot be approximated as a catenoid [28], at least for the membrane tube boundary conditions studied here.

Our model takes explicitly into account lipid flows in the membrane and a drag of the membrane by dynamin moving upon it. Consistent with previous theoretical work [58, 6], constriction of the membrane tube by dynamin involves hydrodynamical flows in the membrane. We find that therefore constriction of tubes covered by long filaments should be characterized by large relaxation times, reaching, for example, about 10 s for membrane shape perturbations of about 250 nm in length. Such slow relaxation suggests that, for in vitro experiments with long membrane tubes, an equilibrium state in the membrane may yet have not been reached when filament fragmentation or membrane fission occurred. On the other hand, for the short, *<* 30 nm, scaffolds proposed to perform scission in vivo [59], membrane flows should equilibrate in under 100 ms, and therefore are unlikely to be rate limiting for membrane scission.

In this study, primary attention has been paid to in vitro experimental setups where long membrane tubes and extended dynamin filaments are typically used, in contrast to short dynamin oligomers involved in endocytocis in living cells. Nonetheless, some conclusions referring to the in vivo situation can be drawn as well. The soft mode and the elasticity of the tetramer should describe the conformational changes taking place in the filament during motor-driven constriction. Dynamin’s large intrinsic twist may help to ensure that non-productive left-handed helices are avoided [60], while low-tension membrane necks compensate by promoting interactions between helical dynamin rings. Later in the process, tension-driven filament expansion may play a role in disassembly of constricted collars. Finally, future studies of the polymer model, extended to include ligand-dependent inter-dimer interactions, will allow hypothetical scenarios of motor function to be tested, and thus contribute towards comprehensive understanding of the dynamin molecular machine.

## Acknowledgments

We are pleased to thank and T. Ando and T. Uchihashi for valuable discussions. J.K.N. and A.S.M. thank Martin Falcke for bringing the subject to their attention. We are grateful to the anonymous referees whose comments have helped us to improve the publication. This work was supported (A.S.M.) by the Ministry of Education, Culture, Sports, Science and Technology of Japan through the World Premier International Research Center Initiative. Financial support (J.K.N.) from the Humboldt Foundation in Germany is gratefully acknowledged. Additional funding support came from the German Research Foundation, SFB740 (project C7 to O.D. and D7 to F.N.).

