## Supplementary Information for "Polymer-like model to study the dynamics of dynamin filaments on deformable membrane tubes"

### Supplementary material for: Polymer-like model to study the dynamics of dynamin filaments on deformable membrane tubes

#### Contents

**1 Locally-cylindrical helix approximation** **7**

**2 Principle component analysis (PCA)** **9**

**3 Repeated translation and rotation defines a helix** **9**

**4 Thermal fluctuations of the curvature and twist** **10**

**5 The weak twist approximation and correlations of the twist angle** **11**

**6 Mapping continuous elastic quantities onto the discretized system** **12**

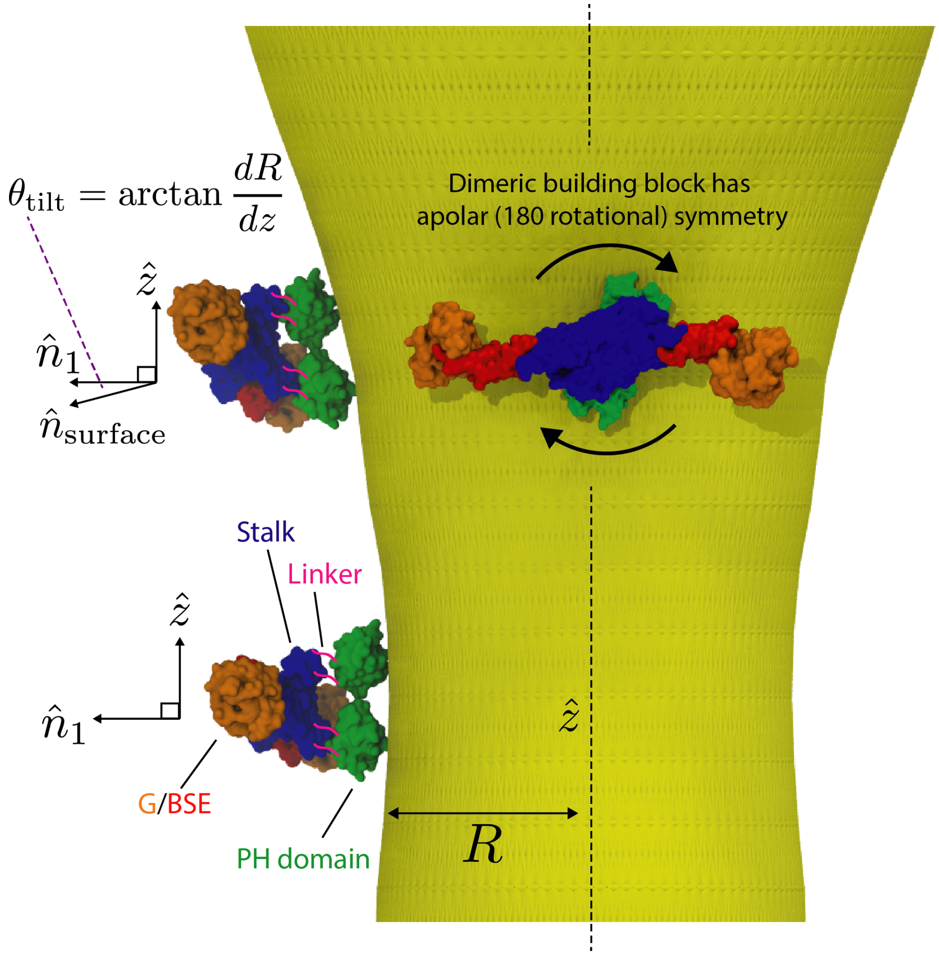

Figure S1: Geometry of the locally-cylindrical helix approximation for the filament. The ribbon alignment vector $\hat{n}_1$ is perpendicular to the membrane tube axis $\hat{z}$. In the non-uniform parts of the tube, this implies a tilt between the PH domains, orthogonal to the membrane, and the stalk domains of the filament. Such tilt has no energy cost because the PH domains are connected via flexible linkers to the stalk. The tilt angle is the angle between the membrane normal vector $\hat{n}_{\rm surface}$ and the filament alignment vector $\hat{n}_1$. Equilibrium filament shapes exhibit the largest radial gradients near filament ends. At low membrane tension $\gamma = 0.01$ $\mathrm{k_BT/nm^2}$, this leads to a maximum tilt of $16^{\rm o}$ reached at the filament end.

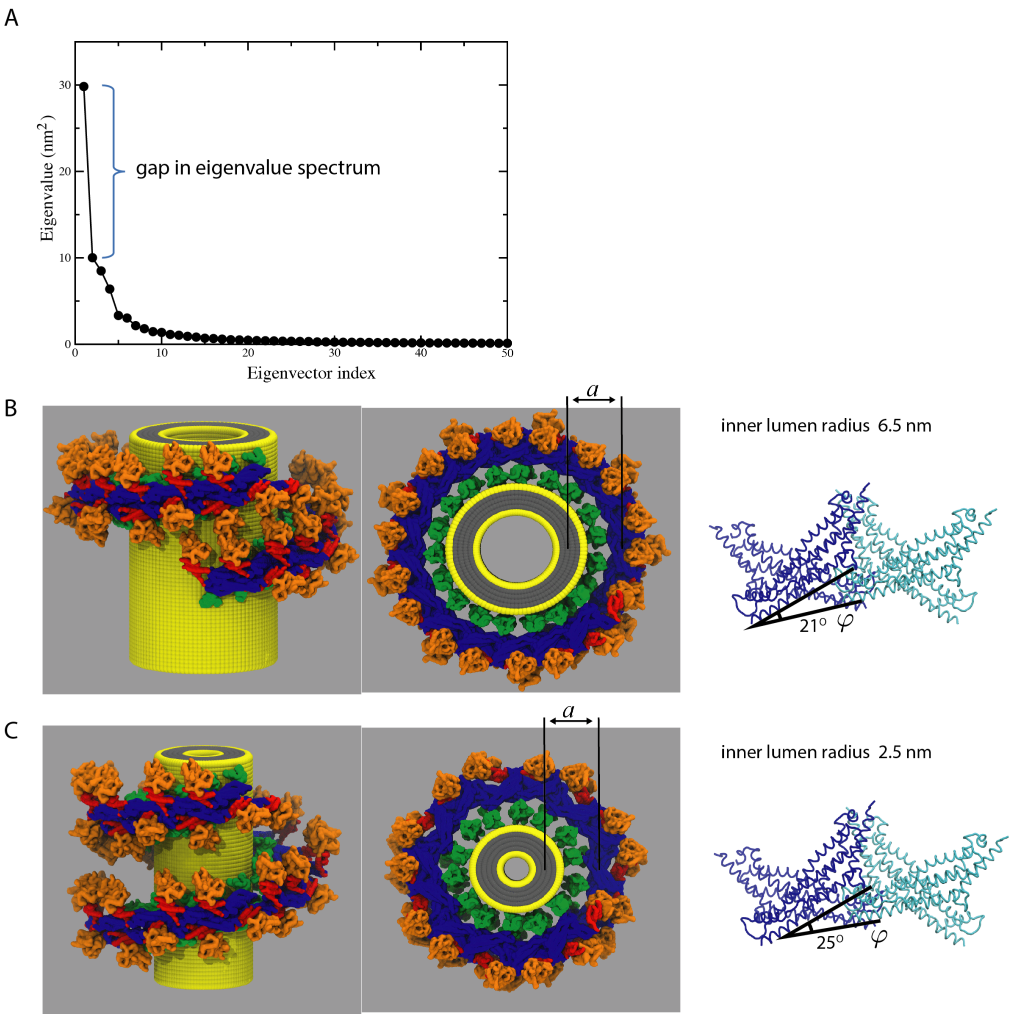

Figure S2: Constricting along the lowest eigenvector in PCA. A) The first 50 eigenvalues from the PCA analysis ordered by size. B) Native tetramer has $\varphi = 21°$ and when many dimers are connected by the native tetramer interface, a filament results with the appropriate dimensions to fit into the non-constricted cryo-EM density [1]. This reconstruction is used to determine $a$, the distance between the center of the filament and the center of the tube bilayer. C) Filament reconstruction for a maximally-compressed tube. Note that the PH domains are connected to the stalk domains by flexible linkers that are not structurally resolved and hence not shown in the figure.

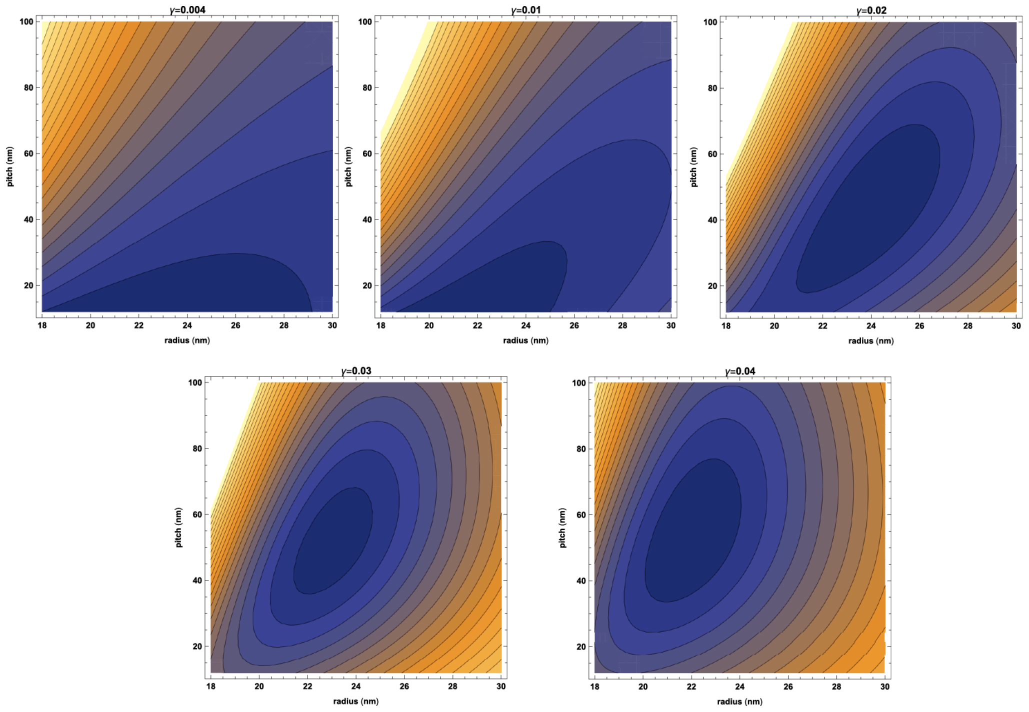

Figure S3: Energy-per-dimer landscapes $g(r, p)$ for different values of the membrane tension $\gamma = 0.004, 0.01, 0.02, 0.03, 0.04$ $\mathrm{k_BT/nm^2}$ and $\chi = 24$ $\mathrm{k_BT}$. These landscapes are determined by Eq. 58 in the main text and using $p = 2\pi h$. The lower bound of the pitch axis is set to the minimum allowed pitch. At low tensions, the energetic minimum is below the minimum pitch, implying maximum compression of the filament. The landscape is relatively narrow with respect to the radius, consistent with the result that the thermal fluctuations are larger for the pitch than the radius (Fig. 9 in the main text). The value of the minimum energy $g_{min}$ per dimer (falling inside the allowed values of radius and pitch) is related to the net cost of deforming the filament/membrane system per unit length. We have $g_{min}$ =6.4, 5.9, 5.1, 4.2, 3.7 $\mathrm{k_BT}$/dimer, respectively, for the various tension values.

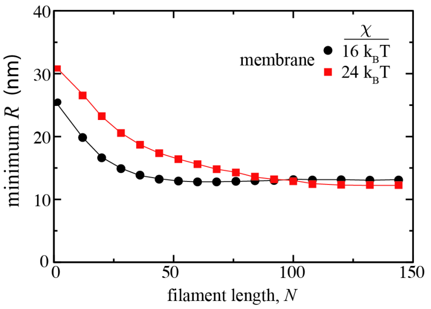

Figure S4: Minimal membrane radius as a function of filament length. The minimum membrane radius achieved along a tube covered by a filament of length $N$ is plotted for two values of membrane stiffness, $\chi = 16$ $\mathrm{k_BT}$(black) and $\chi = 24$ $\mathrm{k_BT}$(red); the membrane tension is $\gamma = 0.0125$ $\mathrm{k_BT/nm^2}$. As the filament grows, it constricts the radius of the underlying tube, while at the same time the deformation of the membrane induces a compression of the filament. Note that the existence of GTP-dependent cross links could result in a small pitch being stabilized for a shorter filament, allowing the minimum radius to be reached for shorter filaments.

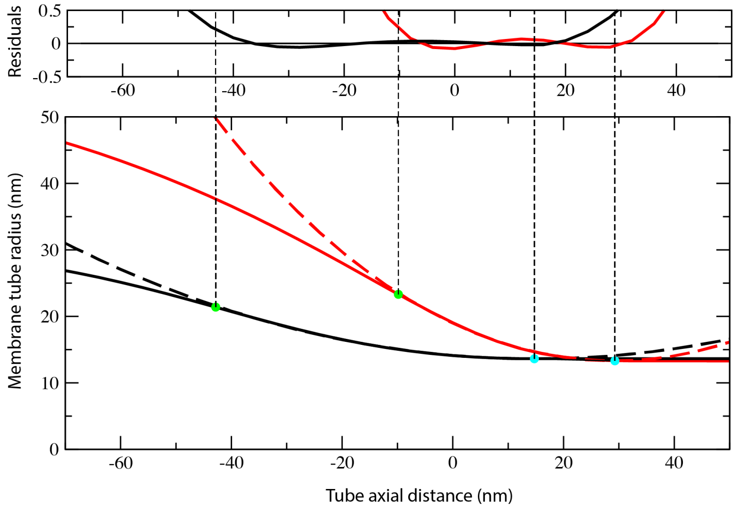

Figure S5: Further examples of the catenary fits to membrane shapes near the end of the filament with membrane stiffness $\chi = 24$ $\mathrm{k_BT}$. Tension is $\gamma = 0.006$ or $0.02$ $\mathrm{k_BT/nm^2}$ for the red and black lines, respectively. Dotted lines show the catenary fitting procedure of Eq. 63 in the main text. For the red curve, $A = 76$ and $z_c = 29$. For the black curve $A = 209$ and $z_c = 15$. The residuals are shown above. The vertical dotted lines are a guide for the eye. The blue dots correspond to $z_c$ and the green dots correspond to the end of the filament.

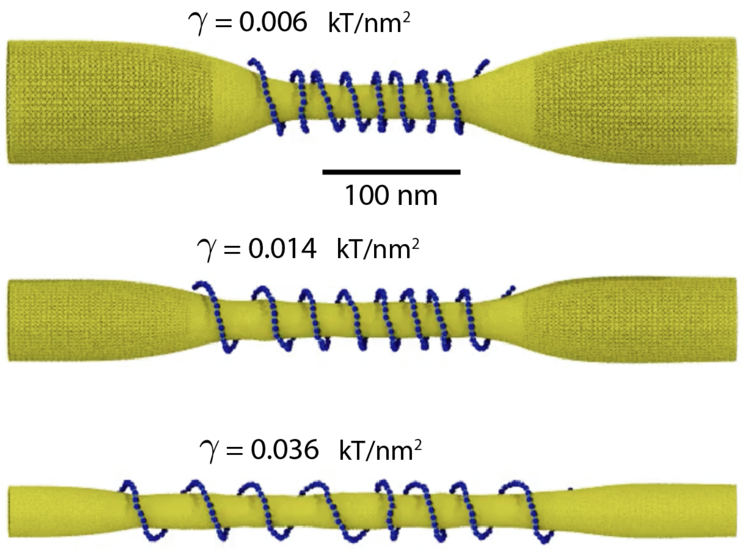

Figure S6: Snapshot from Supplemental Movie 3. Membrane parameters were $\gamma = 0.006, 0.014, 0.036\ \mathrm{k_BT/nm^2}$ and $\chi = 24\ \mathrm{k_BT}$, and the movie corresponds to a small subset of simulations shown in main text Fig. 9. The duration of the movie is 150 ms.

### 1 Locally-cylindrical helix approximation

In this section, the locally-cylindrical helix approximation is derived and discussed. Suppose that the central line of the filament is described in cylindrical coordinates $(\rho, \phi, z)$ by the equations

$$x(\phi) = \rho(z(\phi))\cos\phi, y(\phi) = \rho(z(\phi))\sin\phi, z = z(\phi), \tag{1}$$

where $\rho(z)$ is the dependence of the radius on the axial coordinate. In the locally-cylindrical approximation, it will be assumed that the radius is only slowly varying along the symmetry axis, so that the conditions $(\partial\rho/\partial z)^2) \ll 1$ and $r(\partial^2\rho/\partial z^2) \ll 1$ are satisfied. Below, shorthand notations $\rho_z = \partial\rho/\partial z, \rho_\xi = \partial\rho/\partial\xi$ and $h_\xi = \partial h/\partial\xi$, will be sometimes employed.

The local pitch of the filament is defined as

$$h = \frac{\partial z}{\partial\phi}. \tag{2}$$

If $\xi$ is the arc length along the central line, we have

$$d^2\xi = dz^2 + \rho^2 d\phi^2 + d\rho^2 = (1 + \rho_z^2)dz^2 + \rho^2 d\phi^2 \approx dz^2 + \rho^2 d\phi^2 \tag{3}$$

where we have dropped a small term $\rho_z^2$. Therefore, we approximately find

$$\frac{\partial\phi}{\partial\xi} = \frac{1}{\sqrt{\rho^2 + h^2}} = C \tag{4}$$

where the notation $C = (\rho^2 + h^2)^{-1/2}$ is introduced. The radius and the pitch can be also considered as functions $\rho = \rho(\xi)$ and $h = h(\xi)$ of the intrinsic coordinate $\xi$. Then we approximately have

$$\frac{\partial\rho}{\partial\xi} = \frac{\partial\phi}{\partial\xi}\frac{\partial z}{\partial\phi}\frac{\partial\rho}{\partial z} = Ch\frac{\partial\rho}{\partial z}. \tag{5}$$

Next the orientations of the material frames of the filament should be chosen. Our assumption (not an approximation) is that, for dynamin, the local vectors of the material frame are the same as for a uniform helix, that is

$$\hat{n}_1 = \begin{pmatrix} \cos\phi \\ \sin\phi \\ 0 \end{pmatrix}, \hat{n}_2 = C\begin{pmatrix} h\sin\phi \\ -h\cos\phi \\ \rho \end{pmatrix}, \hat{n}_3 = C\begin{pmatrix} -\rho\sin\phi \\ \rho\cos\phi \\ h \end{pmatrix}. \tag{6}$$

It can be easily checked that these three vectors are mutually orthogonal and have the unit length.

When moving along the filament, the unit vectors of the material frame rotate. The rotation velocities are obtained by taking the derivatives of equations (6). We find

$$\frac{\partial\hat{n}_1}{\partial\xi} = C\begin{pmatrix} -\sin\phi \\ \cos\phi \\ 0 \end{pmatrix}, \tag{7}$$

$$\frac{\partial\hat{n}_2}{\partial\xi} = C\begin{pmatrix} Ch\cos\phi + h_\xi\sin\phi \\ Ch\sin\phi - h_\xi\cos\phi \\ \rho_\xi \end{pmatrix} + \frac{1}{C}\frac{\partial C}{\partial\xi}\hat{n}_2, \tag{8}$$

$$\frac{\partial \hat{n}_3}{\partial \xi} = C \begin{pmatrix} -C\rho\cos\phi - \rho_\xi \sin\phi \\ -C\rho\sin\phi + \rho_\xi\cos\phi \\ h_\xi \end{pmatrix} + \frac{1}{C}\frac{\partial C}{\partial \xi}\hat{n}_3. \tag{9}$$

The above equations can be used to determine the two curvatures and the twist:

$$\kappa = \frac{\partial \hat{n}_3}{\partial \xi}\hat{n}_1 = C^2\rho = \frac{\rho}{\rho^2+h^2}, \tag{10}$$

$$\tau = -\frac{\partial \hat{n}_1}{\partial \xi}\hat{n}_2 = C^2 h = \frac{h}{\rho^2+h^2}, \tag{11}$$

$$\sigma = \frac{\partial \hat{n}_3}{\partial \xi}\hat{n}_2 = C^2(\rho h_\xi - h\rho_\xi) = \kappa\frac{\partial h}{\partial \xi} - \tau\frac{\partial \rho}{\partial \xi}. \tag{12}$$

By using the inverse relationships

$$\rho = \frac{\kappa}{\kappa^2+\tau^2}, h = \frac{\tau}{\kappa^2+\tau^2}, \tag{13}$$

the last equation for the geodesic curvature can be also written as

$$\sigma = \rho\frac{\partial \tau}{\partial \xi} - h\frac{\partial \kappa}{\partial \xi}. \tag{14}$$

Moreover, the geodesic curvature can also be expressed in terms of the twist angle. Because $\tan\theta = \tau/\kappa$, we obtain

$$\sigma = \frac{\partial \theta}{\partial \xi}. \tag{15}$$

It should be noted that, if a filament is attached to a membrane, its radius is fixed at $\rho = R + a$ by the local radius $R$ of the membrane tube. Therefore, the validity condition $(\partial\rho/\partial z)^2 \ll 1$ of the locally-cylindrical helix approximation for the filament is equivalent to the validity condition $(\partial R/\partial z)^2 \ll 1$ of the locally-cylindrical approximation for the membrane.

The locally-cylindrical helix approximation can also be applied if different assumptions about the orientations of local material frames for the filament are made. Below, we briefly show the results for the case when the filament is assumed to be always orthogonal to the membrane surface, i.e. if there is no tilt.

In this case, the central line of the filament is still described by equations (1), but the local vectors of the material frame are different from those given by equations (6). Instead, we approximately have

$$\hat{n}_1 = \begin{pmatrix} \cos\phi \\ \sin\phi \\ \rho_z \end{pmatrix}, \hat{n}_2 = C\begin{pmatrix} h\sin\phi + \rho\rho_z\cos\phi \\ -h\cos\phi + \rho\rho_z\sin\phi \\ \rho \end{pmatrix}, \hat{n}_3 = C\begin{pmatrix} -\rho\sin\phi + \rho_z h\cos\phi \\ \rho\cos\phi + \rho_z h\sin\phi \\ h \end{pmatrix}. \tag{16}$$

In these equations, we have already dropped the terms of order $\rho_z^2$ or higher. Using Eqs. 16, the velocity vectors $\partial\hat{n}_{1,2,3}/\partial\xi$ can be again found, allowing the determination of the curvatures and the twist. We omit the derivation and present the final approximate results: The normal curvature and the twist are still given by Eqs. 10 and 11, but the expression for the geodesic curvature acquires a correction term,

$$\sigma = \kappa \frac{\partial h}{\partial \xi} - \tau \frac{\partial \rho}{\partial \xi} - \frac{1}{h} \frac{\partial \rho}{\partial \xi}. \quad (17)$$

In the derivation of these results, we had to assume, in addition to $\rho_z^2 \ll 1$, that the condition $\rho\rho_{zz} \ll 1$ is also satisfied. In the the expressions for $\kappa, \tau$ and $\sigma$, the terms of the order $\rho_z^2$ and $\rho\rho_{zz}$ or higher are dropped.

In principle, the complete analysis in the manuscript could have been repeated also for the filaments without a tilt, i.e. always being orthogonal to the membrane surface. For that, only a correction term for the geodesic curvature in the expression for the elastic energy of the filament would have to be introduced.

### 2 Principle component analysis (PCA)

To determine the large amplitude correlated motions, we computed the principle components from the explicit solvent molecular dynamics trajectory of the stalk tetramer. The GROMACS tool `g_covar` was used for the analysis [2].

The lowest eigenmode (largest eigenvector) corresponds to the largest amplitude deformation mode. The significant gap in the eigenvalue spectrum suggests that the lowest mode is a good estimate for how the filament responds to stress (Fig. S2A). Supplemental Movie 1 shows the effect of moving along the lowest eigenvector. The tetramer interface is relatively more flexible than the dimer interface. For each frame in the movie, the deformed tetramer interface is used to identically connect 18 crystal dimers, which creates a filament. As the tetramer is moved along the lowest eigenvector, the collective effect is to reduce the radius of the filament as the angle $\varphi$ between each dimer increases. Concomitantly, the twist increases and results in a slight increase in pitch. The first and last frames of the movie are shown in Figure S2. $\varphi$ changing by only four degrees is sufficient to allow the filament to adapt from an inner lumen radius of 6.5 nm (non-constricted state [1]) to and inner lumen radius of 2.5 nm, small enough that spontaneous hemifusion should occur. The next three largest eigenvectors mostly affect the pitch, and include small rearrangements within the packing of the coiled-coil in the monomer. These rearrangements are likely artifacts of having only one tetramer interface explicitly represented, since the changes in structure only happen away from the tetramer interface.

### 3 Repeated translation and rotation defines a helix

Given a shape conserving transformation (translation $\vec{T}$ and rotation matrix $\mathbf{R}$) between monomers (monomer here means repeating unit; for the dynamin helix, the repeating unit is a dimer) in a helical filament, the shape of the helix that is created when the transformation is repeatly applied can be determined (Fig. S7). The vector $\vec{a}_{i,i+1} = \vec{r}_i - \vec{r}_{i+1}$ connecting the centers of mass of monomers $i$ and $i+1$ transforms to the vector connecting the $i+1$ and $i+2$ monomers by

$$\vec{a}_{i+1,i+2} \rightarrow \mathbf{R}(\vec{a}_{i,i+1} + \vec{T}) \equiv \Theta. \quad (18)$$

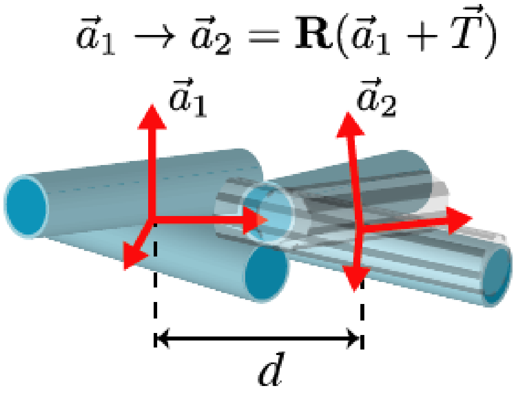

Figure S7: Cartoon describing the fitting procedure of the stalk tetramer simulation trajectory. For each snapshot in the trajectory (stalk domain represented by cyan cylinder), crystal dimers (gray outline) are fit to the tetramer. The transformation between them determines the separation distance $d$, curvature $\kappa$, and twist $\tau$ corresponding to its tetrameric interface.

$|\vec{T}| = |\vec{a}_{i,i+1}|$ so it is simplest to take $\vec{T} = \vec{a}_{i,i+1}$. Note that $|\vec{T}| = d$, the stretching coordinate. Any three dimensional rotation can be described by an axis $\vec{u}$ (a vector which is unchanged by the rotation) and an angle $\theta$ of rotation about $\vec{u}$. Since $\mathbf{R}\vec{u} = \mathbf{I}\vec{u}$, $\vec{u}$ is seen as an eigenvector of $\mathbf{R}$ with eigenvalue $\lambda = 1$. By determining the eigenvectors of $\mathbf{R}$, $\vec{u}$ is determined. In this context, $\vec{u}$ is the axis of the helix. To determine $\theta$, take a vector $\vec{v}$ perpendicular to $\vec{u}$ and calculate the angle between $\vec{v}$ and $\mathbf{R}\vec{v}$. A simpler way is the identity $\mathrm{Tr}\mathbf{R} = 1 + 2\cos\theta$. Now, armed with $\vec{u}$ and $\theta$, one can calculate $r$ and $p$, the radius and pitch of the helix. The idea is that displacement along $\vec{u}$ is related to $p$ and perpendicular displacement is related to $r$. It can be shown that

$$r = |\vec{u} \times \vec{T}| \frac{\cos\theta/2}{\sin\theta}. \tag{19}$$

and

$$h = \frac{\vec{u} \cdot \vec{T}}{2\pi\theta} \tag{20}$$

$h$ is the mathematical pitch; the pitch that is measured experimentally as the separation distance between successive turns is $2\pi h$. Thus, given any arbitrary translation and rotation, it will always form a homogeneous helix when repeatedly applied. The curvature $\kappa$ and twist $\tau$ for the helix can be calculated using the identities:

$$\kappa = \frac{r}{r^2 + p^2}, \quad \tau = \frac{p}{r^2 + p^2}. \tag{21}$$

### 4 Thermal fluctuations of the curvature and twist

In main text Section 3.1, thermal fluctuations of curvature and twist are used to determine the elastic parameters of the filament, $\kappa_0, \tau_0, \alpha_\kappa$ and $\alpha_\tau$. Here, we derive analytical expressions for such fluctuations in the continuum approximation. Because our MD simulations do not include the membrane, we shall estimate the fluctuations neglecting the energy contributions from the membrane, i.e. for a free dynamic filament.

Generally, thermal fluctuations obey the Boltzmann distribution

$$P = Z^{-1} \exp(-\frac{\delta E}{k_B T}) \tag{22}$$

where $\delta E$ is the energy associated with them, $T$ is the temperature, $k_B$ is the Boltzmann constant, and $Z$ is a normalization factor.

Because membrane elasticity is not involved, only a change in the elastic energy of the filament has to be estimated. Since thermal fluctuations are relatively weak, we can introduce local deviations of the curvature and the twist from their equilibrium values, $\kappa = \kappa_0 + \delta\kappa$ and $\tau = \tau_0 + \delta\tau$ and keep only second-order terms in such deviations in $\delta E$. This yields

$$\delta E_{\mathrm{F}} = \frac{1}{2}\int_0^L [\beta(\rho_0 \frac{\partial \delta\tau}{\partial \xi} - h_0 \frac{\partial \delta\kappa}{\partial \xi})^2 + \alpha_\kappa \delta\kappa^2 + \alpha_\tau \delta\tau^2] d\xi. \tag{23}$$

Suppose that we have chosen a short filament element of length $\Delta L$ with curvature $\delta\kappa$ and twist $\delta\tau$. Since the curvature and the twist do not vary within the element, the first term in $\delta E_{\mathrm{F}}$ vanishes and

$$\delta E_{\mathrm{F}} = \frac{1}{2}\alpha_\kappa \delta\kappa^2 \Delta L + \frac{1}{2}\alpha_\tau \delta\tau^2 \Delta L. \tag{24}$$

Therefore, local curvature and twist obey Gaussian distributions

$$P(\kappa) = \frac{1}{\sqrt{2\pi\sigma_\kappa}} \exp[-\frac{(\kappa - \kappa_0)^2}{\sigma_\kappa}] \tag{25}$$

$$P(\tau) = \frac{1}{\sqrt{2\pi\sigma_\tau}} \exp[-\frac{(\tau - \tau_0)^2}{\sigma_\tau}] \tag{26}$$

with the dispersions

$$\sigma_\kappa = \frac{2k_B T}{\alpha_\kappa \Delta L}, \sigma_\tau = \frac{2k_B T}{\alpha_\tau \Delta L}. \tag{27}$$

### 5 The weak twist approximation and correlations of the twist angle

A different derivation needs to be performed to estimate the elastic modulus $\beta$, the elasticity modulus for geodesic curvature fluctuations. As we show below, this parameter controls the correlation length of the twist angle $\theta$ in a special situation when the filament is wrapped around a stiff rod. To determine such correlation length, we further assume that the filament is *weakly* twisted, i.e., the condition $\theta \ll 1$ or, equivalently, $h \ll \rho$, holds. The weak twist approximation is not used in the rest of our study. It can be checked that it is actually violated at the free ends of the filament where the local twist may become large (main text Fig. 7B). On the other hand, it is applicable in the middle, uniform part of a compressed filament. Indeed, if the pitch is $p$ =12 nm and the filament radius is $\rho$ =20 nm, we have $h = p/2\pi$=2.9 nm and the twist angle is $\theta = 0.095$.

If $h \ll \rho$, the continuum description becomes simplified. In this limit,

$$\theta = \frac{h}{\rho}, \kappa = \frac{1}{\rho}, \tau = \frac{h}{\rho^2} = \frac{\theta}{\rho}. \tag{28}$$

In this case, the elastic energy of the filament is approximately

$$E_{\rm F} = \frac{1}{2}\int_0^L [\beta(\frac{\partial\theta}{\partial\xi})^2 + \alpha_\kappa(\frac{1}{\rho} - \kappa_0)^2 + \alpha_\tau(\frac{\theta}{\rho} - \tau_0)^2 +]d\xi \tag{29}$$

If the filament is wrapped around a rigid rod and its radius is fixed at $\rho_0$, we have (dropping a constant term)

$$E_{\rm F} = \frac{1}{2}\int_0^L [\beta(\frac{\partial\theta}{\partial\xi})^2 + \frac{\alpha_\tau}{\rho_0^2}(\theta - \theta_0)^2]d\xi \tag{30}$$

where $\theta_0 = \rho_0\tau_0$. Thus, the filament will have the equilibrium twist angle $\theta = \theta_0$. Decomposing $\theta(\xi)$ into a superposition of plane waves,

$$\theta(\xi) = \theta_0 + \sum_k \theta_k e^{ik\xi}, \tag{31}$$

the energy can be written furthermore as

$$E_{\rm F} = \frac{1}{2}\sum_k (\frac{\alpha_\tau}{\rho_0^2} + \beta k^2) |\theta_k|^2 . \tag{32}$$

In the equilibrium Boltzmann distribution, each normal mode should bear, on the average, the same energy $\mathrm{k_B T}$. This implies that

$$\left\langle |\theta_k|^2 \right\rangle = \frac{2k_B T}{(\alpha_\tau/\rho_0^2) + \beta k^2}. \tag{33}$$

The correlation function

$$C(\xi) = \langle(\theta(\xi) - \theta_0)(\theta(0) - \theta_0)\rangle \tag{34}$$

of twist angle fluctuations can be written as

$$C(\xi) = \sum_k \left\langle |\theta_k|^2 \right\rangle e^{ik\xi}. \tag{35}$$

Substituting $\left\langle |\theta_k|^2 \right\rangle$ in this equation and performing the summation, one finds that

$$C(\xi) \sim \exp(-|\xi|/l_\theta) \tag{36}$$

and the correlation length $l_\theta$ for fluctuations of the twist angle is

$$l_\theta = \rho_0\sqrt{\frac{\beta}{\alpha_\tau}}. \tag{37}$$

In Section 3.3, Eq. 37 is used to estimate the elastic modulus $\beta$ from the coarse-grained simulations.

### 6 Mapping continuous elastic quantities onto the discretized system

This section explicitly writes out the derivatives of the energy used to calculate forces in the polymer model.

#### 6.1 Filament

The elastic ribbon is mapped onto a chain of beads with coordinates ($r_i$,$\phi_i$,$z_i$) that are harmonically constrained to their nearest neighbors. The terms that contribute to the filament energy in terms of $\xi$:

$$E_{\mathrm{F}} = E^{\kappa} + E^{\tau} + E^{\sigma} + E^{\lambda} + E^{\mathrm{stretch}} \tag{38}$$

$$E_{\mathrm{F}} = \frac{1}{2}\int_{-\mathcal{L}/2}^{\mathcal{L}/2} \left[\alpha_\kappa(\kappa-\kappa_0)^2 + \alpha_\tau(\tau-\tau_0)^2 + \beta\sigma^2 + \lambda(\rho - R(z(\xi)) - a)^2\right] d\xi + E^{\mathrm{stretch}} \tag{39}$$

$$\kappa = \frac{\rho}{\rho^2+h^2},\ \tau = \frac{h}{\rho^2+h^2},\ \sigma = \rho\frac{d\tau}{d\xi} - h\frac{d\kappa}{d\xi},\ h = \frac{dz}{d\phi} \tag{40}$$

Forces on the beads $i$ come from discretizing along $\xi$ in intervals of $d_0$.

$$\kappa_i = \frac{\rho_i}{\rho_i^2+h_i^2},\ \tau_i = \frac{h_i}{\rho_i^2+h_i^2},\ \sigma_i = \rho_i\frac{d\tau_i}{d\xi} - h_i\frac{d\kappa_i}{d\xi},\ h_i = \frac{z_{i+1}-z_i}{\phi_{i+1}-\phi_i} = \frac{\Delta z_i}{\Delta\phi_i},\ \frac{dX_i}{d\xi} = \frac{X_i - X_{i-1}}{d_0} \tag{41}$$

$h$ is defined in terms of a ratio of differences of coordinates so the partial derivatives are useful:

$$\frac{\partial h_j}{\partial z_i}\begin{cases}\frac{-1}{\Delta\phi_i}, & \text{if } j = i\\ \frac{1}{\Delta\phi_i}, & \text{if } j = i-1\end{cases} \tag{42}$$

and

$$\frac{\partial h_j}{\partial \phi_i}\begin{cases}\frac{\Delta z_i}{(\Delta\phi_i)^2}, & \text{if } j = i\\ \frac{-\Delta z_i}{(\Delta\phi_i)^2}, & \text{if } j = i-1\end{cases} \tag{43}$$

The forces for each energetic term are divided into an expression per coordinate. Some useful expressions:

$$\Delta\kappa_i = \frac{\rho_i}{\rho_i^2+h_i^2} - \kappa_0,\ \Delta\tau_i = \frac{h_i}{\rho_i^2+h_i^2} - \tau_0 \tag{44}$$

$$A_i = \frac{\rho_i^2 - h_i^2}{(\rho_i^2+h_i^2)^2},\ B_i = \frac{-2h_i\rho_i}{(\rho_i^2+h_i^2)^2} \tag{45}$$

For bending:

$$F^{\kappa}_{\rho_i} = -\frac{\delta E^{\kappa}}{\delta\rho_i} = -\alpha_\kappa\Delta\kappa_i(-A_i) \tag{46}$$

$$\rho_i F^{\kappa}_{\phi_i} = -\frac{\delta E^{\kappa}}{\delta\phi_i} = -\alpha_\kappa\Delta\kappa_j B_j\frac{dh_j}{d\phi_i} = -\alpha_\kappa\left[\Delta\kappa_i B_i\frac{\Delta z_i}{(\Delta\phi_i)^2} - \Delta\kappa_{i-1}B_{i-1}\frac{\Delta z_{i-1}}{(\Delta\phi_{i-1})^2}\right] \tag{47}$$

$$F^{\kappa}_{z_i} = -\frac{\delta E^{\kappa}}{\delta z_i} = -\alpha_\kappa\Delta\kappa_j B_j\frac{dh_j}{dz_i} = -\alpha_\kappa\left[-\Delta\kappa_i B_i\frac{1}{\Delta\phi_i} + \Delta\kappa_{i-1}B_{i-1}\frac{1}{\Delta\phi_{i-1}}\right] \tag{48}$$

For the twist:

$$F^{\tau}_{\rho_i} = -\frac{\delta E^{\tau}}{\delta\rho_i} = -\alpha_\tau\Delta\tau_i B_i \tag{49}$$

$$\rho_i F^{\tau}_{\phi_i} = -\frac{\delta E^{\tau}}{\delta\phi_i} = -\alpha_\tau\Delta\tau_j A_j\frac{dh_j}{d\phi_i} = -\alpha_\tau\left[\Delta\tau_i B_i\frac{\Delta z_i}{(\Delta\phi_i)^2} - \Delta\tau_{i-1}B_{i-1}\frac{\Delta z_{i-1}}{(\Delta\phi_{i-1})^2}\right] \tag{50}$$

$$F^{\tau}_{z_i} = -\frac{\delta E^{\tau}}{\delta z_i} = -\alpha_\tau\Delta\tau_j A_j\frac{dh_j}{dz_i} = -\alpha_\tau\left[-\Delta\tau_i B_i\frac{1}{\Delta\phi_i} + \Delta\tau_{i-1}B_{i-1}\frac{1}{\Delta\phi_{i-1}}\right] \tag{51}$$

For the geodesic curvature:

$$F^{\sigma}_{\rho_i} = -\frac{\delta E^{\sigma}}{\delta \rho_i} = -\beta\sigma_i \left[\tfrac{d}{d\xi}\tau_i + \rho_i \tfrac{d}{d\xi}B_i - h_i \tfrac{d}{d\xi}(-A_i)\right] \tag{52}$$

$$\rho_i F^{\sigma}_{\phi_i} = -\frac{\delta E^{\sigma}}{\delta \phi_i} = -\beta\left[\sigma_j\rho_j \tfrac{d}{d\xi}A_j \tfrac{dh_j}{d\phi_i} - \sigma_j \tfrac{dh_j}{d\phi_i}\tfrac{d}{d\xi}\kappa_j - \sigma_j h_j \tfrac{d}{d\xi}B_j \tfrac{dh_j}{d\phi_i}\right] \tag{53}$$

$$= -\beta\left(\sigma_i\rho_i \tfrac{d}{d\xi}A_i \tfrac{\Delta z_i}{(\Delta\phi_i)^2} - \sigma_{i-1}\rho_{i-1}\tfrac{d}{d\xi}A_{i-1}\tfrac{\Delta z_{i-1}}{(\Delta\phi_{i-1})^2}\right) \tag{54}$$

$$+ \beta\left(\sigma_i \tfrac{\Delta z_i}{(\Delta\phi_i)^2}\tfrac{d}{d\xi}\kappa_i - \sigma_{i-1}\tfrac{\Delta z_{i-1}}{(\Delta\phi_{i-1})^2}\tfrac{d}{d\xi}\kappa_{i-1}\right) \tag{55}$$

$$+ \beta\left(\sigma_i h_i \tfrac{d}{d\xi}B_i \tfrac{\Delta z_i}{(\Delta\phi_i)^2} - \sigma_{i-1}h_{i-1}\tfrac{d}{d\xi}B_{i-1}\tfrac{\Delta z_{i-1}}{(\Delta\phi_{i-1})^2}\right) \tag{56}$$

$$F^{\sigma}_{z_i} = -\frac{\delta E^{\sigma}}{\delta z_i} = -\beta\left[\sigma_j\rho_j \tfrac{d}{d\xi}A_j \tfrac{dh_j}{dz_i} - \sigma_j \tfrac{dh_j}{dz_i}\tfrac{d}{d\xi}\kappa_j - \sigma_j h_j \tfrac{d}{d\xi}B_j \tfrac{dh_j}{dz_i}\right] \tag{57}$$

$$= -\beta\left(-\sigma_i\rho_i \tfrac{d}{d\xi}A_i \tfrac{1}{\Delta\phi_i} + \sigma_{i-1}\rho_{i-1}\tfrac{d}{d\xi}A_{i-1}\tfrac{1}{\Delta\phi_{i-1}}\right) \tag{58}$$

$$+ \beta\left(-\sigma_i \tfrac{1}{\Delta\phi_i}\tfrac{d}{d\xi}\kappa_i + \sigma_{i-1}\tfrac{1}{\Delta\phi_{i-1}}\tfrac{d}{d\xi}\kappa_{i-1}\right) \tag{59}$$

$$+ \beta\left(-\sigma_i h_i \tfrac{d}{d\xi}B_i \tfrac{1}{\Delta\phi_i} + \sigma_{i-1}h_{i-1}\tfrac{d}{d\xi}B_{i-1}\tfrac{1}{\Delta\phi_{i-1}}\right) \tag{60}$$

For the interaction with the membrane:

$$F^{\lambda}_{\rho_i} = -\frac{\delta E^{\lambda}}{\delta \rho_i} = -\lambda(\rho_i - R_i(z(\xi)) - a) \tag{61}$$

$$\rho_i F^{\lambda}_{\phi_i} = -\frac{\delta E^{\lambda}}{\delta \phi_i} = 0 \tag{62}$$

$$F^{\lambda}_{z_i} = -\frac{\delta E^{\lambda}}{\delta z_i} = -\lambda(\rho_i - R_i(z(\xi)) - a)\tfrac{dR_i(z(\xi))}{dz} \tag{63}$$

where $R_i$ means the radius of the membrane disk belonging to bead $i$. Several beads can share a disk. $\frac{dR_i(z)}{dz}$ denotes the membrane gradient measured at the disk belonging to bead $i$. This gradient is expressed in the next section. The stretching potential is a standard nearest-neighbor harmonic bond:

$$E^{\text{stretch}} = \alpha_{\text{stretch}}\left(|\vec{r}_{i+1} - \vec{r}_i| - d_0\right)^2 \tag{64}$$

#### 6.2 Membrane

The membrane is represented by $M$ cylindrical disks, each of which has height $d_{\text{M}}$ and is centered along the axis $R = 0$. The $j$th disk is fixed at $z_j$, but free to vary its radius $R_j$. There are two principle curvatures $c_1$ and $c_2$. $c_1$ is simply the inverse radius, i.e. $c_{1,j} = 1/R_j$. The second is the curvature in an orthogonal direction, along $z$, and thus can be approximated as the second derivative of the radius along $z$, $c_{2,j} \approx \frac{d^2 R_j}{dz^2}$. For a continuous cylindrically symmetric membrane of length $L$ with a dynamin filament of length $\mathcal{L}$ covering the membrane from $z_1$ to $z_2$, the energy per unit length along $z$ is:

$$E_{\rm M} = E^\gamma + E^\chi + E^\lambda \tag{65}$$

$$E_{\rm M} = \pi \int_0^L \left[2\gamma R + \tfrac{\chi}{R} + \chi R \left(\tfrac{\partial^2 R}{\partial z^2}\right)^2\right] dz + \tfrac{1}{2}\lambda \int_0^{\mathcal{L}} (\rho(\xi) - R(z(\xi)) - a)^2 d\xi \tag{66}$$

$$= \pi \int_0^L \left[2\gamma R + \tfrac{\chi}{R} + \chi R \left(\tfrac{\partial^2 R}{\partial z^2}\right)^2\right] dz + \tfrac{1}{2}\lambda \int_{z_1}^{z_2} (r(\xi(z)) - R(\xi(z)) - a)^2 \tfrac{\partial \xi}{\partial z} dz \tag{67}$$

$$\tag{68}$$

Determining the surface stress $\sigma_{\rm m}$ on disk $j$ requires a functional derivative

$$2\pi\sigma_{\rm m} = -\frac{\delta E_{\rm M}}{\delta R_j(z)} = -\left[\frac{\partial E}{\partial R_j(z)} + \partial_z^2 \cdot \frac{\partial E}{\partial R_j''(z)}\right] \tag{69}$$

For the tension:

$$2\pi\sigma_{\rm m}^\gamma = -\frac{\delta E^\gamma}{\delta R_j} = -2\pi\gamma \tag{70}$$

For the bending:

$$2\pi\sigma_{\rm m}^\chi = -\frac{\delta E^\chi}{\delta R_j} = \pi\chi\left[-\tfrac{1}{R_j^2} + \left(\tfrac{\partial^2 R_j}{\partial z^2}\right)^2 + \tfrac{\partial^2}{\partial z^2}\left(2R_j \tfrac{\partial^2 R_j}{\partial z^2}\right)\right] \tag{71}$$

$$= \pi\chi\left[-\tfrac{1}{R_j^2} + 3\left(\tfrac{\partial^2 R_j}{\partial z^2}\right)^2 + 4\tfrac{\partial R_j}{\partial z}\tfrac{\partial^3 R_j}{\partial z^3} + 2R_j \tfrac{\partial^4 R_j}{\partial z^4}\right] \tag{72}$$

The derivatives along the membrane profile can be symmetrically discretized along the membrane disks:

$$\frac{\partial R_j}{\partial z} = \frac{R_{j+1} - R_{j-1}}{2d_{\rm M}} \tag{73}$$

$$\frac{\partial^2 R_j}{\partial z^2} = \frac{R_{j+1} - 2R_j + R_{j-1}}{d_{\rm M}^2} \tag{74}$$

$$\frac{\partial^3 R_j}{\partial z^3} = \frac{R_{j+2} - 2R_{j+1} + 2R_{j-1} - R_{j-2}}{2d_{\rm M}^3} \tag{75}$$

$$\frac{\partial^4 R_j}{\partial z^4} = \frac{R_{j+2} - 4R_{j+1} + 6R_j - 4R_{j-1} + R_{j-2}}{d_{\rm M}^4} \tag{76}$$

$$\tag{77}$$

In writing the Helfrich energy of the membrane (Eq. 66), a cross term of the form $\frac{1}{R}\left(\frac{\partial^2 R}{\partial z^2}\right)$, the so-called "Gaussian curvature," was neglected. The functional derivative of the membrane energy explicitly shows that a Gaussian curvature term gives no contribution to the force because of the cancellation of one of the normal curvatures ($1/R$) and the integration area element ($2\pi R$):

$$\left[\frac{\partial}{\partial R(z)} + \partial_z^2 \cdot \frac{\partial}{\partial R''(z)}\right] \cdot R\frac{1}{R}\left(\frac{\partial^2 R}{\partial z^2}\right) = 0 \tag{78}$$

#### 6.3 Interaction between the filament and the membrane

For the interaction with the filament:

$$2\pi\sigma_{\mathrm{m}}^{\lambda} = -\frac{\delta E^{\lambda}}{\delta R_j} = \frac{\sum_{i\in j}\lambda(\rho_i - R_j - a)\frac{\Delta z_i}{d_0}}{N_j} \tag{79}$$

The sum over $i \in j$ means all beads $i$ that belong to disk $j$ and $N_j$ is the total number of beads $i$ that belong to disk $j$. $\frac{\Delta\xi_i}{\Delta z_i} = d_0/\Delta z_i$ gives the conversion between length along the filament and length along the membrane. The interaction term gives the average force per unit $z$ provided by the filament. Eq. 79 is only correct if the local pitch is never negative (left-handed), which can happen under thermal fluctuations. A better form is to calculate $\theta_j$, the average twist angle for the filament interacting with membrane disk $j$ which is simply the total length of the filament interacting with $j$ divided by the height of a membrane disk $d_{\mathrm{M}}$, $\theta_j = N_j d_0/d_{\mathrm{M}}$. The stress from the interaction with the filament then becomes:

$$2\pi\sigma_{\mathrm{m}}^{\lambda} = -\frac{\delta E^{\lambda}}{\delta R_j} = \frac{d_0}{d_M}\sum_{i\in j}\lambda(\rho_i - R_j - a) \tag{80}$$
